# Antimicrobials from a feline commensal bacterium inhibit skin infection by drug-resistant *S. pseudintermedius*

**DOI:** 10.1101/2021.03.08.432323

**Authors:** Alan M. O’Neill, Kate A. Worthing, Nikhil Kulkarni, Fengwu Li, Teruaki Nakatsuji, Robert H. Mills, Joyce Y. Cheng, David J. Gonzalez, Jacqueline M. Norris, Richard L. Gallo

**Affiliations:** Department of Dermatology, University of California, San Diego, CA, USA; College of Veterinary Medicine, University of Arizona, Tucson, USA; Department of Pharmacology, University of California, San Diego, CA, USA; Skaggs School of Pharmacy and Pharmaceutical Sciences, University of California, San Diego; Sydney School of Veterinary Science, University of Sydney, NSW, Australia

## Abstract

Methicillin-resistant *Staphylococcus pseudintermedius* (MRSP) is an important emerging zoonotic pathogen that causes severe skin infections. To combat infections from drug-resistant bacteria, the transplantation of commensal antimicrobial bacteria as a therapeutic has shown clinical promise. We screened a collection of diverse staphylococcus species from domestic dogs and cats for antimicrobial activity against MRSP. A unique strain (*S. felis* C4) was isolated from feline skin that inhibited MRSP and multiple gram-positive pathogens. Competition experiments in mice showed that *S. felis* significantly reduced MRSP skin colonization and an antimicrobial extract from *S. felis* significantly reduced necrotic skin injury from MRSP infection. Fluorescence and electron microscopy revealed that *S. felis* antimicrobials disrupted bacterial but not eukaryotic cell membranes. LC/MS identified several *S. felis* phenol-soluble modulin beta (PSMβ) peptides that exhibited antimicrobial and anti-inflammatory activity. These findings indicate a feline commensal bacterium that could be utilized in bacteriotherapy against difficult-to-treat animal and human skin infections.

## INTRODUCTION

Skin is colonized by hundreds of diverse bacterial species that exist within a complex and dynamic chemical landscape. These chemical interactions can play important roles in skin health, immune education and protection against pathogen colonization and infection (Sanford and Gallo, 2013). The composition of the skin microbial community of humans and animals varies extensively, in part due to different skin habitats, i.e. increased hair density in animals, as well as more subtle differences in the chemical and biological conditions of the skin (Grice and Segre, 2011; Ross et al., 2018). Overall, the human microbial skin community is distinct from and significantly less diverse than that of both wild and domestic animals (Ross et al., 2018). Human skin is generally dominated by few taxa present at high abundance e.g. cutibacteria, streptococci and staphylococci, whereas canine skin harbors a more equally distributed and diverse group of taxa (Song et al., 2013). Naturally, close contact between humans and animals can be a source for microbial transmission (Frana et al., 2013; Lai et al., 2017). Although it remains to be determined if shared taxa are stable over time, there are reports that exposure to pets early in life can be protective against atopic disease in later life (Mandhane et al., 2009).

In contrast, there are also many documented cases of human staphylococcal infection from epidemiological exposure to dogs (Somayaji et al., 2016). Companion animals can act as reservoirs for methicillin-resistant *S. aureus* (MRSA) and more commonly, *S. pseudintermedius* (MRSP), with both species sharing many common invasion and virulence factors (Garbacz et al., 2013). The zoonotic significance of *S. pseudintermedius* may have been previously underestimated because it was frequently misidentified as *S. aureus* in human wound infections (Börjesson et al., 2015). More advanced diagnostic techniques such as MALDI-TOF have led to increased detection of human *S. pseudintermedius* infections (Ference et al., 2019). Colonization of *S. pseudintermedius* is a contributing factor in canine atopic dermatitis (AD). Interestingly, the prevalence of AD in humans and AD in dogs are similar (10-15% in US) and present with remarkably similar immunological and clinical manifestations (Marsella and Girolomoni, 2009; Silverberg, 2019). Likewise, several studies have reported a decrease in the microbiome diversity of AD and increased colonization of *S aureus* in humans and *S. pseudintermedius* in dogs (Fazakerley et al., 2009; Nakatsuji and Gallo, 2019; Older et al., 2020). In human AD, *S. aureus* was identified in higher relative abundances during disease flares (Kong et al., 2012). Similarly, the relative abundance of *S. pseudintermedius* was also shown to increase with disease flares in canine AD (Bradley et al., 2016). Common treatment modalities exist for both diseases. Dilute bleach baths are a common antiseptic treatment for AD, with the goal of reducing the carriage of staphylococci (Banovic et al., 2018; Chopra et al., 2017). However, its effectiveness as an antibacterial agent is controversial (Sawada et al., 2019).

An alternative and promising approach is not to disrupt but to re-establish a healthy microbiome community on the skin. To do this our group and others have screened for naturally occurring, and sometimes rare commensal species on healthy human skin that express antimicrobial activity against skin pathogens. A recent example is the discovery of commensal staphylococcus strains that produce lantibiotics that when applied to skin of patients with AD, demonstrated clinically improved symptoms and reductions of lesional *S. aureus* counts (Nakatsuji et al., 2017; Nakatsuji et al. in press). Other studies have reported antimicrobial staphylococcus strains belonging to species of *S. lugdunensis* (Zipperer et al., 2016), *S. epidermidis* (Cogen et al., 2010) and *S. capitis* (O’Neill et al., 2020) isolated from different niches of the skin and nares. In contrast, very little is known regarding the antimicrobial activity of animal-derived staphylococci and their clinical potential against skin infection. Here we identified *S. felis* C4, a potent antimicrobial isolate from feline skin that inhibited the growth of MRSP *in vitro* and *in vivo. S. felis* C4 produced several α-helical amphipathic peptides that demonstrated antimicrobial and anti-inflammatory activity. This discovery represents a potential new bacteriotherapeutic for human and canine skin diseases associated with *S. pseudintermedius* colonization and infection.

## RESULTS

### A screen of animal-derived staphylococci isolates identifies a feline skin commensal bacterium with broad-spectrum antimicrobial activity

We sought to determine whether commensal staphylococci collected from the skin, nasal, oral and perineal sites of companion dogs and cats exhibit antimicrobial activity against methicillin-resistant *S. pseudintermedius* (MRSP) ST71 (Fig. 1A) (Ma et al., 2020; K. Worthing et al., 2018). Fifty-eight staphylococcus isolates across the coagulase-positive (CoPS) and coagulase-negative (CoNS) groups were screened, including validated antimicrobial strains of human origin, *S. hominis* A9 (Nakatsuji et al., 2017) and *S. capitis* E12 (O’Neill et al., 2020) and a non-active *S. aureus* 113 negative control strain (Fig. 1B). The animal test isolates were screened for antimicrobial activity by live co-culture on agar plates or in the presence of sterile conditioned supernatant, as illustrated in Fig. 1A. Amongst all test isolates, five strains demonstrated greater than 80% inhibition of *S. pseudintermedius* growth (dashed line) across all three different dilutions of supernatant (1:1, 1:4, 1:8) (Fig. 1C). Surprisingly, these strains exhibited greater potency compared to the positive control *S. hominis* A9 supernatant (indicated by black circle), which inhibited growth of *S. pseudintermedius* in a 1:1 dilution, but not at 1:4 or lower. Amongst the five positive hits, three were identified as *S. felis* and two *S. pseudintermedius*. In the second independent antimicrobial assay, all five isolates including positive control *S. hominis* A9, produced an observable zone of inhibition against *S. pseudintermedius* during live co-culture on agar (Fig. 1D). The two feline *S. felis* species (C4, N26 labelled with white arrows) produced the largest inhibitory zones, extending 3.0-3.3 mm outward from the edge of the growing colony. The *S. felis* C4 strain was chosen for further analysis, as it demonstrated potent activity and was isolated from healthy skin. To investigate the significance and selectivity of the *S. felis* antimicrobial supernatant, we tested its capacity to inhibit the growth of other clinically relevant, gram-positive and gram-negative pathogens (of which several belong to the clinically-relevant ESKAPE group). Of the four gram-negative strains tested, only moderate inhibition was demonstrated after 18 h incubation with 80-100% undiluted *S. felis* C4 supernatant (Fig. 1E). In contrast, culture with just 1-5% of *S. felis* C4 supernatant was sufficient to inhibit >80% growth of all four gram-positive organisms, including *S. pseudintermedius, E. faecium, B. subtilis* and *S. aureus*.

**Figure 1.**
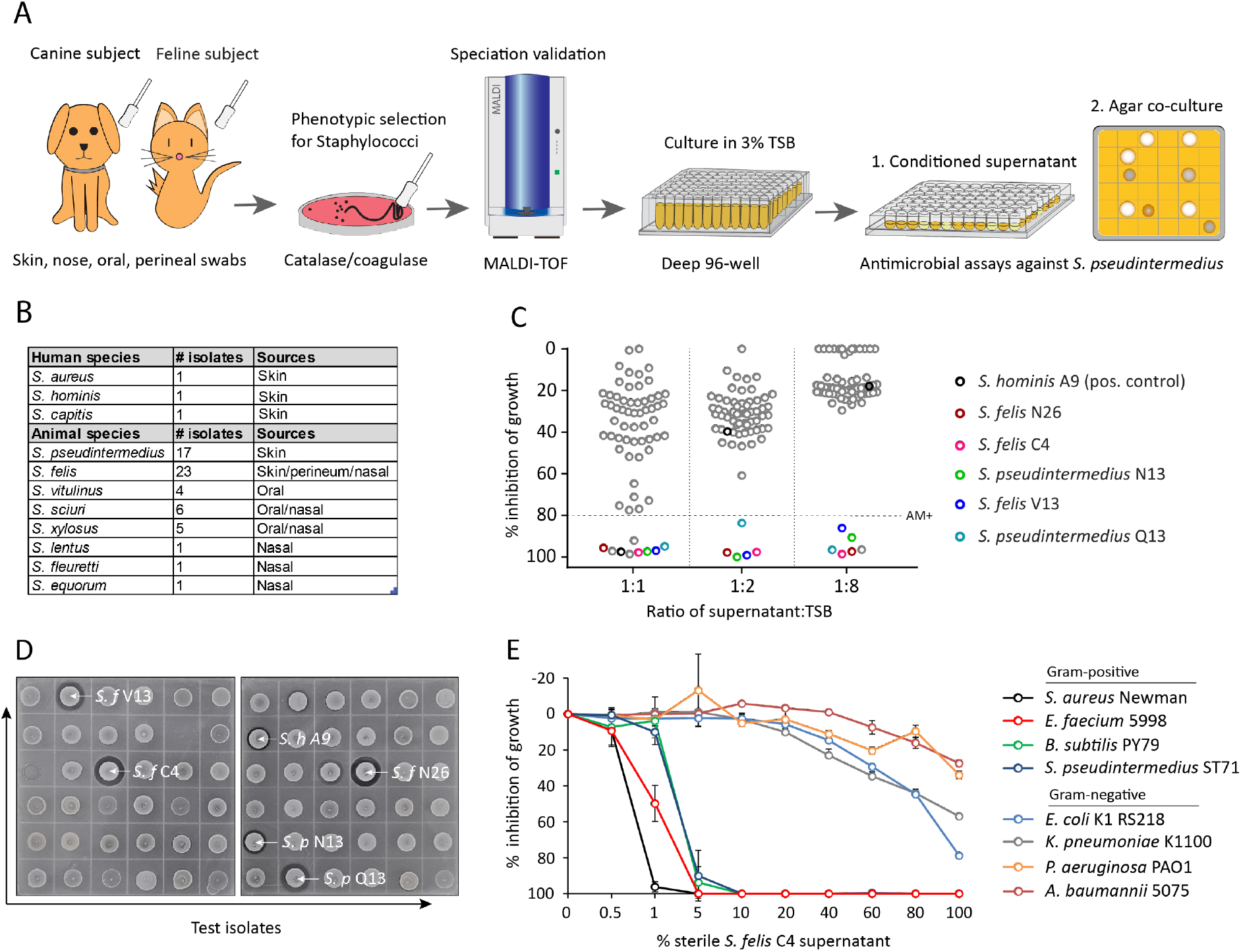
Screening and discovery of a feline skin commensal bacterium that inhibits drug-resistant gram-positive pathogens. **(A)** Illustration of the selection and screening strategy of animal-derived staphylococci against the growth of methicillin-resistant *S. pseudintermedius* (MRSP) ST71 in liquid culture and agar co-culture assays. **(B)** The panel of 58 feline and canine isolates selected for antimicrobial testing, including human-derived *S. hominis* A9 and *S. capitis* E12 positive control antimicrobial strains and the non-antimicrobial *S. aureus* 113 negative control. **(C)** Inhibition of *S. pseudintermedius* ST71 growth by OD600, relative to TSB control at 100%, after 18 h incubation in 50%, 25%, or 12.5% (1:1, 1:4, 1:8 ratio) sterile conditioned supernatant from all staphylococci isolates. Greater than 80% inhibition of growth was considered antimicrobial (AM+). **(D)** Images of the agar co-culture assay showing zone of inhibition (black circle surrounding colony) produced by all staphylococci test isolates against *S. pseudintermedius* ST71, including *S. felis* C4, N26, V13 (*S. f* C4, *S. f* N26, *S. f* V13), *S. pseudintermedius* N13 and Q13 (*S. p* N13 and *S. p* Q13), and positive control *S. hominis* A9 (*S. h* A9, all indicated by arrows). **(E)** Inhibition of bacterial growth relative to TSB alone at 100%, against select gram-positive and gram-negative pathogens after 18 h incubation (48 h incubation for *E. faecium*) in the presence of increasing amounts of sterile conditioned supernatant from *S. felis* C4 overnight growth. Error bars indicate SEM. Representative of three separate experiments.

Of the three antimicrobial *S. felis* isolates, only the C4 supernatant retained activity after precipitation with 75% ammonium sulfate (AS) (Suppl. Fig. 1A). Moreover, 75% AS was highly effective in precipitating the antimicrobial factor(s) from the C4 supernatant since no activity could be visualized in the non-precipitate fraction (Suppl. Fig. 1B). This effect was also achieved with a simpler extraction by n-butanol. The antimicrobial butanol extract remained active at room temperature (RT), up to one week and was stable after boiling (Suppl. Fig. 1C). As expected, the butanol extraction provided a partially purified and enriched antimicrobial fraction compared to crude supernatant and AS precipitation (Suppl. Fig 1D). Therefore, further experiments involving the *S. felis* C4 extract describes sterile supernatant obtained via n-butanol extraction. Interestingly, the antimicrobial activity of *S. felis* C4 was sufficient to disrupt bacterial biofilms. Reports have shown that most clinically-derived *S. pseudintermedius* strains are biofilm producers (Singh et al., 2013). Biofilm formation is considered an important determinant of staphylococci virulence and is associated with increased skin colonization and severity of disease (Di Domenico et al., 2018; Kwiecinski et al., 2015). A 4 h preformed biofilm of*pseudintermedius* ST71 showed a significant decrease in crystal violet (CV) staining over time, when incubated with 100% conditioned supernatant of *S. felis* C4, indicating biofilm disruption and degradation (Suppl. Fig. 2A). Importantly, the *S. felis* C4 extract also exhibited similar anti-biofilm activity as crude conditioned supernatant, with biofilm mass reduced by 48% at 250 μg/ml and 58% at 500 μg/ml (Suppl. Fig. 2B).

### *S. felis* C4 inhibits *S. pseudintermedius* skin colonization and infection in mice

We next sought to investigate the translational potential of *S. felis* C4 and its antimicrobial products as a potential therapy against *S. pseudintermedius* colonization and infection. Since *S. felis* C4 is a commensal bacterium that was isolated from healthy feline skin, we speculated it should be safe and well tolerated on mouse skin. *S. felis* C4 was found to be sensitive to several common antibiotics (Fig. 2A) and as such represents a suitable strain for further investigation as a bacteriotherapy. We therefore assessed the skin tolerability of a 3-day topical application of *S. felis* C4 on SKH1 hairless mice. Whereas *S. pseudintermedius* and the non-antimicrobial *S. felis* ATCC 49168 induced evidence of scaling and redness on mouse back skin, *S. felis* C4 did not promote any adverse reaction (Fig. 2B).

**Figure 2.**
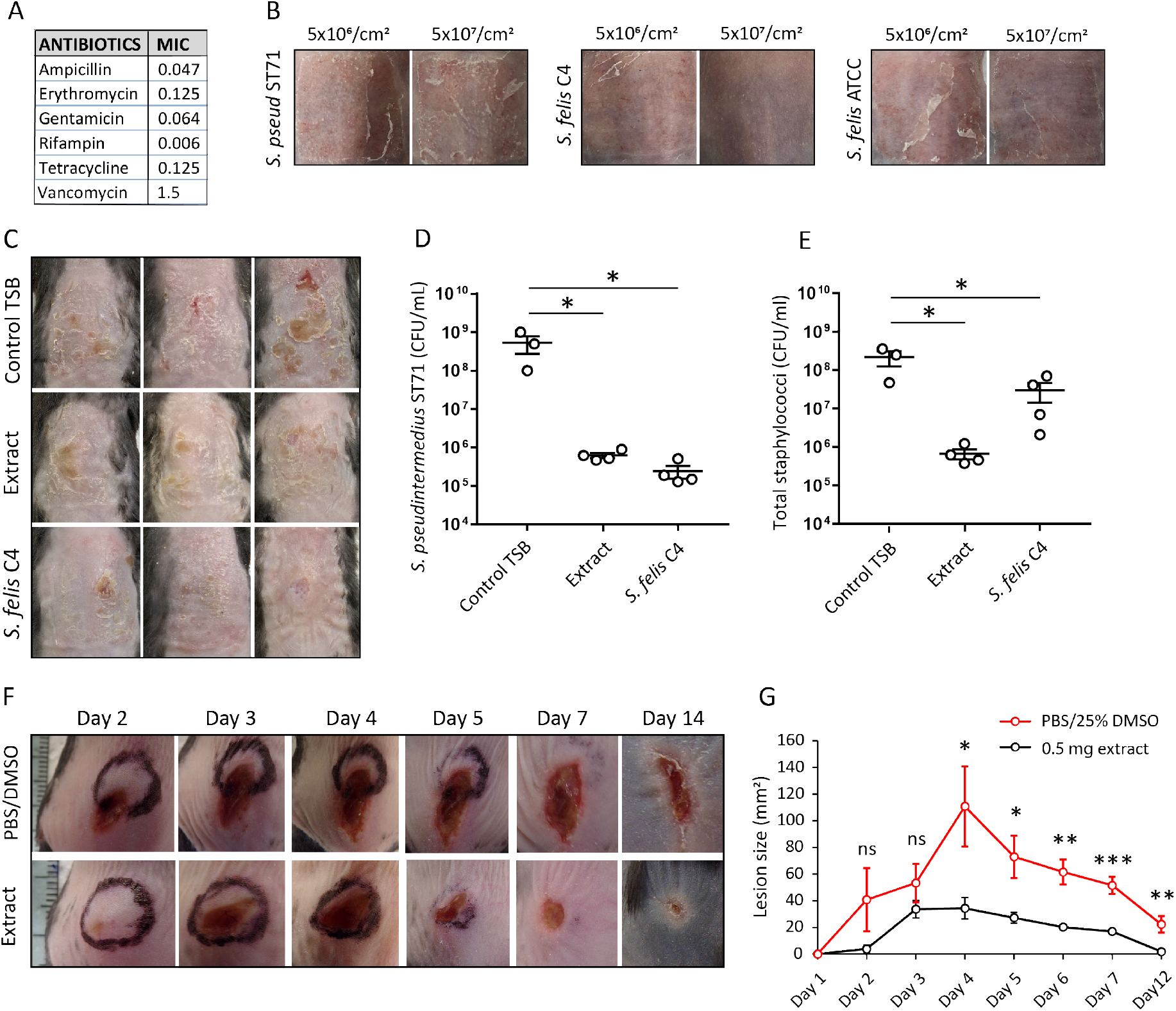
Live bacteriotherapeutic intervention with *S. felis* C4 protects against *S. pseudintermedius* colonization in mice. **(A)** Minimum inhibitory concentrations (MIC) of the indicated antibiotics against *S. felis* C4. **(B)** Representative images of the dorsal skin of 8-10 week-old SKH1 mice 3 days post-challenge with live *S. felis* C4, *S. pseudintermedius* ST71 (*S. pseud* ST71) or *S. felis* ATCC 49168, inoculated at the indicated amounts. n=2, per treatment group **(C-E)** 5 x 10^7^ CFU/cm^2^ of *S. pseudintermedius* ST71 was applied onto the back skin of C57BL/6 mice for 48 h and challenged with TSB, *S. felis* C4 extract (1 mg) or live *S. felis* C4 (5 x 10^7^ CFU/cm^2^) for 72 h. Post-treatment, mouse back skin was photographed **(C)** and swabbed to enumerate *S. pseudintermedius* ST71 CFU on selective Baird-Parker egg yolk tellurite agar **(D)** or total staphylococci CFU on selective mannitol-salt agar plates **(E)**. n=3 for TSB and n=4 for extract and *S. felis* C4. Error bars indicate SEM. One-way ANOVA with multiple comparisons (Dunnett’s correction) was performed. *p* values: \**p* < 0.05; **(F-G)** At day 0, 1 x10^7^ CFU of *S. pseudintermedius* ST71 was intradermally injected into the back skin of 8-10 week-old C57BL/6 mice and at 1 h post-infection two inoculations of *S. felis* C4 extract (250 μg) or PBS/25% DMSO control were injected adjacent to the infection site. **(F)** Representative images of *S. pseudintermedius* ST71-induced dermonecrosis over time after receiving control PBS/DMSO or *S. felis* C4 extract. **(G)** Quantification of lesion size (mm^2^) over time as measured by L x W of lesions. n=4 for DMSO/PBS and n=5 for extract. Error bars indicate SEM. A two-tailed, unpaired Student’s *t*-test was performed. *p* values: \**p* < 0.05; \*\**p* < 0.01; \*\*\**p* < 0.001.

To test the antimicrobial activity of *S. felis* C4 on skin, we applied 5 x 10^7^ CFU/cm^2^ *S. pseudintermedius* directly onto mouse back skin for 48 h, then applied an equal density of *S. felis* C4 or 1 mg of extract to the infected site. This was repeated daily for three days. The back skin showed a reduction in scaling and redness post-treatment with *S. felis* C4 or extract compared to control (Fig. 2C). Enumeration of *S. pseudintermedius* ST71 CFUs after plating onto selective agar revealed a significant 2.9 log decrease in CFU after extract application and a 3.3 log decrease after live *S. felis* C4 application (Fig. 2D). Enumeration of total staphylococci CFUs after plating onto selective agar once again confirmed that the extract treatment significantly reduced bacterial colonization (Fig. 2E). In contrast, total staphylococci CFU counts after the *S. felis* C4 application were more similar to the control group, suggesting that the *S. felis* bacteria successfully colonized and outcompeted *S. pseudintermedius* ST71 on the skin (Fig. 2E). To further test *S. felis* C4 extract as an anti-MRSP intervention, we evaluated its efficacy in limiting the infectious outcome of cutaneous challenge with *S. pseudintermedius*. An inoculum of 1 x 10^7^ CFU *S. pseudintermedius* ST71 was intradermally administered into the back skin of mice. At 1 h post-infection, two intradermal inoculations of 250 μg extract were administered adjacent to the infection site and necrosis was monitored by measuring lesion size over a 14-day period. Compared to controls, the extract-treated mice exhibited slower lesion progression from day 1 to 2, and significantly better protection from *S. pseudintermedius* skin disease from day 4 (Fig. 2F-G). These results demonstrate the *in vivo* efficacy and clinical potential for *S. felis* C4 as a bacteriotherapy against *S. pseudintermedius* skin colonization and infection.

### Antimicrobial *S. felis* C4 promotes disruption of the bacterial membrane

Next, we sought to better understand how the antimicrobial action of *S. felis* C4 negatively affects bacterial physiology. Given the selective nature of the *S. felis* C4 supernatant against gram-positives (Fig. 1E), we speculated that *S. felis* C4 activity may target and compromise the bacterial membrane and/or cell wall. To address this question, transmission electron microscopy (TEM) imaging was conducted on sectioned *S. pseudintermedius* ST71 bacteria exposed to 1 h treatment with control DMSO (1%), sub-MIC (1 μg/ml), MIC (8 μg/ml) or 5X MIC (40 μg/ml) extract concentrations (Fig. 3A). TEM observations upon control DMSO treatment showed normal, uniform spherical cocci physiology and septum formation indicating active replication. In contrast, short exposure to the extract resulted in drastic changes to the bacterial ultrastructural morphology, with evidence of cell wall thickening and alterations in the structure and rigidity of the cell membrane (Fig. 3A, lower zoom inset panels). Moreover, treatment with extract showed evidence of greater chromosomal compaction compared to control, evidenced by the increased (electron) density of the nucleoid (highlighted yellow arrows). Additional evidence supporting a role for *S. felis* C4 in promoting bacterial membrane damage was provided by several *in vitro* microbial cell viability assays. From mid-log phase cultures of *S. pseudintermedius* ST71 30 min post-treatment with the extract, a dose-dependent increase in reactive oxygen species (ROS) (Fig. 3B) coincided with a concomitant decrease in intracellular ATP levels (Fig. 3C). Further evidence for a membrane-active antimicrobial was provided by increased dual staining of SYTO9-positive bacteria (green) with the membrane-impermeable dye propidium iodide (PI) (red). After treatment with 5X MIC of extract, a proportion of bacterial cells showed dual staining, with entry of PI likely reflecting a loss of membrane integrity (Fig. 3D).

**Figure 3.**
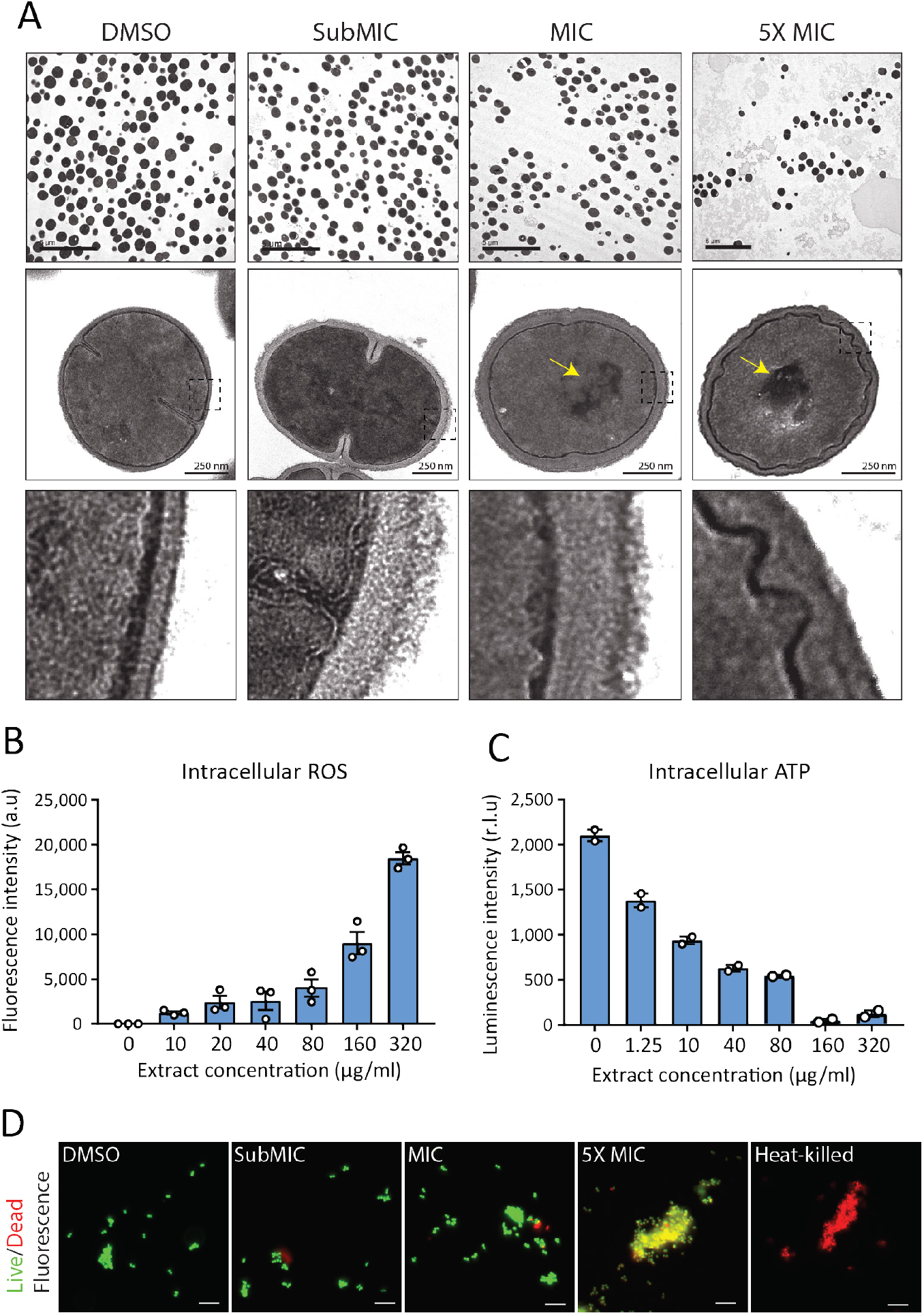
*S. felis* C4 extract contains a membrane-active antibacterial molecule. (**A**) TEM images of *S. pseudintermedius* ST71 after 1 h treatment with DMSO control or *S. felis* C4 extract at indicated concentrations. Yellow arrows indicate evidence of condensed DNA. Lower image panels represent higher magnification of regions highlighted by dashed black boxes. Scale = 250 nm (**B**) Total ROS accumulation in *S. pseudintermedius* ST71 after 1 h treatment in increasing concentrations of *S. felis* C4 extract. Error bars indicate SEM. (**C**) Total intracellular ATP accumulation in *S. pseudintermedius* ST71 after 1 h treatment in increasing concentrations of *S. felis* C4 extract. Error bars indicate SEM. (**D**) Live/dead fluorescent images of *S. pseudintermedius* ST71 after 1 h treatment with DMSO control or extract at indicated concentrations. Viable bacterial cells were stained green by SYTO9 and damaged/dead cells were stained red by PI. Scale = 10 μm. (**B**), (**C**) representative of two separate experiments.

### Purification and identification of PSMβ peptides as antimicrobial products of *S. felis* C4

To determine the nature of the antimicrobial product produced by *S. felis* C4, sterile supernatant was purified by HPLC. This yielded two major peaks that eluted at 44% and 47% acetonitrile (Suppl. Fig. 3A). Anti-*S. pseudintermedius* activity was predominantly associated with fraction 32 which eluted at 55% acetonitrile (Suppl. Fig. 3B). SDS PAGE and silver stain of fraction 32 and neighboring inactive fractions revealed a unique band of roughly 5 kDa in size (Suppl. Fig. 3C). To determine if this small protein was responsible for antimicrobial activity, gel slices of the fraction 32 lane corresponding to small, medium and larger proteins (≤5 kDa, 5-20 kDa and 20-50 kDa, respectively) were excised and extracted by acetone precipitation, as previously described (Botelho et al., 2010; Zhang et al., 2015). Only the ≤5 kDa band demonstrated antimicrobial activity after incubation with *S. pseudintermedius* (Suppl. Fig. 3D), thereby suggesting the likely candidate to be a small peptide. Mass spectrometry (MS) analysis of the top 8 hits in the active and non-active fractions identified several putative small antimicrobial peptides (AMP), representing the phenol soluble modulin beta (PSMβ1-3) and gamma (PSMγ, aka delta-hemolysin) families and a peptide of unknown function containing a EF-hand domain, common amongst some antimicrobial Ca^2+^ binding proteins, such as S100A8/S100A9 (Chazin, 2011) (Suppl. Fig. 3E). Whole genome analysis of the *S. felis* C4 strain confirmed the presence of three PSMβ-encoding genes (Suppl. Fig. 3F). Like some mammalian cationic AMPs such as cathelicidin LL-37, the PSMβ and EF-hand domain peptides have an α-helical amphipathic-like formation, a structural motif conserved in the recognition and binding of biological membranes (Suppl. Fig. 3G). To determine if the peptides identified by MS exhibited antimicrobial activity, synthetic versions of all three *S. felis* PSMβ1-3 and the EF-hand domain containing peptide were tested. Interestingly, all three PSMβ peptides inhibited *S. pseudintermedius* growth at a concentration of 50 μg/ml (Fig. 4A). In contrast, the EF-hand domain-containing peptide did not inhibit bacterial growth up to 200 μg/ml. These results suggest that PSMβ peptides could be mediating the antimicrobial activity of *S. felis* C4.

**Figure 4.**
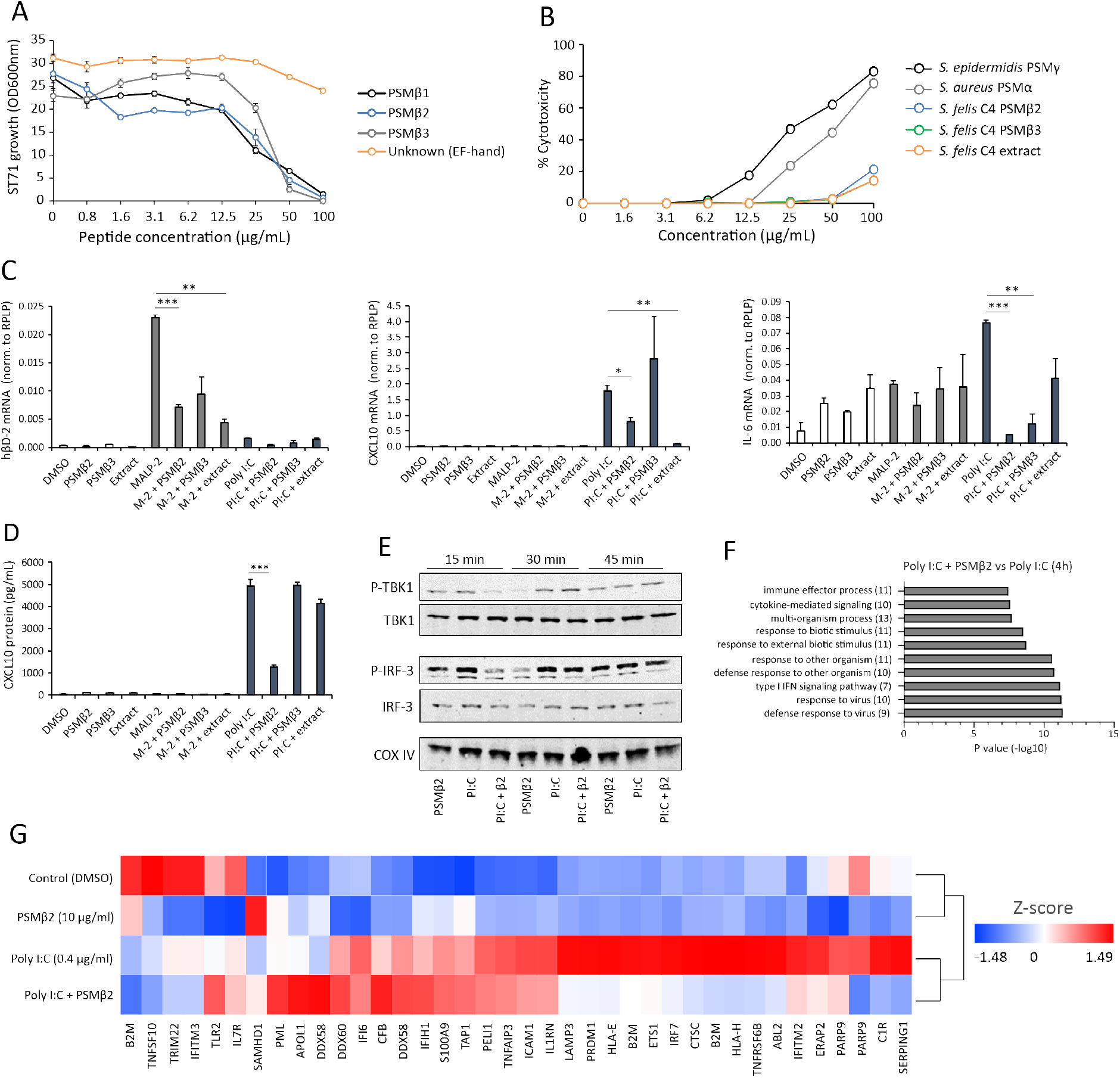
Antimicrobial *S. felis* C4 extract and PSMβ suppress TLR-mediated inflammation. **(A)** Growth of *S. pseudintermedius* ST71 after 18 h incubation with increasing concentrations of synthetic formylated PSMβ peptides (PSMβ1, PSMβ2, PSMβ3), or synthetic peptide containing an EF-hand domain (Unknown EF-hand). **(B)** Lactate dehydrogenase (LDH) release in NHEKs after 24 h treatment with *S. felis* C4 extract, *S. felis* PSMβ2, PSMβ3 or positive control cytotoxic PSMα from *S. aureus* or PSMγ from *S. epidermidis*. Percentage (%) cytotoxicity measured by maximum LDH release into supernatant collected after untreated cell freeze thaw. **(C)** mRNA transcript abundance measured by qPCR in NHEKs stimulated with or without TLR2/6 agonist MALP-2 (200 ng/ml) or TLR3 agonist Poly I:C (0.4 μg/ml) in the presence or absence of *S. felis* C4 extract, PSMβ2 or PSMβ3 (10 μg/ml) or DMSO control (0.1%) at 4 h post-treatment. **(D)** Quantification of CXCL10 protein by ELISA from supernatant of NHEKs stimulated with MALP-2 or Poly I:C in the presence of *S. felis* extract or PSMβ2, PSMβ3 or DMSO control 24 h post-treatment. **(E)** Time-course of the TLR3 signaling cascade by immunoblot of phosphorylated TBK1 and IRF3 proteins after stimulation of NHEKs with Poly I:C, DMSO, or co-treatment with Poly I:C and PSMβ2. **(F)** Gene ontology (GO) pathway analysis of genes downregulated in NHEKs after 4 h co-treatment with Poly I:C and PSMβ2 versus treatment with Poly I:C alone. **(G)** Hierarchical clustering and Heatmap visualization of selected genes from GO enriched ‘immune response’ pathway (1.5-fold change) 4 h post-treatment with DMSO, PSMβ2 or Poly I:C alone or with PolyI:C and PSMβ2 co-treatment. (**A-D**) error bars indicate SEM. Representative of at least three separate experiments. One-way ANOVA with multiple corrections (Turkey correction) was performed. *p* values: * p< 0.05; ** p<0.01; *** p<0.001.

### *S. felis* C4 extract and PSMβ peptides exhibit anti-inflammatory activity by suppressing TLR-mediated inflammation

Unlike the well characterized cytolytic and inflammatory activities of PSMα, a defined role for PSMβ in mediating host interactions has been largely unexplored (Da et al., 2017). Based on our previous observation that *S. felis* C4 was well tolerated on murine skin, we asked if the antimicrobial extract and individual PSMβs would be tolerated by human keratinocytes. Primary normal human keratinocytes (NHEK) were treated with increasing concentrations of different PSMs for 24 h and cytotoxicity determined by quantifying LDH release. As expected, the hemolytic toxins PSMα and PSMγ were found to be highly cytotoxic whereas *S. felis* PSMβ2, PSMβ3, and the extract resulted in less than 5% LDH release at the antimicrobial concentration of 50 μg/ml (Fig. 4B).

To investigate the potential effects of *S. felis* C4 on the host immune response, we stimulated cells with various TLR agonists in the presence or absence of the extract or the individual PSMs and measured inflammatory gene expression. NHEKs were treated with *S. felis* PSMβ2, PSMβ3, extract, DMSO control alone, or each in combination with the TLR2 agonist MALP-2 (200 ng/μl) or the TLR3 agonist Poly I:C (0.4 μg/ml) for 4 h. Neither PSMβ nor the extract resulted in any detectable increase in gene expression, whereas MALP-2 significantly increased the expression of hBD-2, and Poly I:C significantly increased CXCL10 and IL-6 expression (Fig. 4C). Interestingly, these TLR-mediated responses were significantly reduced during co-treatment with PSMβ or extract. This result was confirmed by ELISA, showing that PSMβ2 had a significant effect on suppressing CXCL10 secretion in NHEKs after 24 h cotreatment (Fig. 4D). To determine if this interaction is specific for epithelial cells, we also stimulated human THP-1 macrophage-like cells with MALP-2, or the TLR4 agonist LPS and found that IL-6 and TNFα expression was decreased during co-treatment with PSMβ2 (Suppl. Fig. 4). The addition of PSMβ2 to NHEK activated by poly I:C demonstrated that PSMβ2 inhibited phosphorylation of TBK1 and IRF3 at 15 min post-stimulation with the peptide (Fig. 4E). This inhibition of inflammatory target gene and kinase activity was further evaluated by the analysis of changes in global gene expression using RNA-Seq analysis of NHEKs stimulated with Poly I:C, with and without PSMβ2 at 4 h and 24 h. Gene ontology (GO) analyses revealed the significant down-regulation of several gene clusters associated with ‘immune effector process’ and ‘type I IFN signaling’ at 4 h during co-treatment of PSMβ2 and Poly I:C (Fig. 4F). A heatmap of selected genes within the ‘Immune response’ GO term at 4 h post-treatment further highlighted the suppressive effect, but more significantly, showed that PSMβ2 treatment alone did not induce an immunological response in NHEKs (Fig. 4G). However, when NHEKs were treated with PSMβ2 in the absence of an inflammatory stimulus, we identified the downregulation of genes within GO terms such as “response to interleukin-1” (Suppl. Fig. 5A) and “pathogenic E. coli infection” (Suppl. Fig. 5B), suggesting that exposure to PSMβ2 primes cells to dampen potential inflammatory mediators in response to bacterial ligands. This contrasted with PSMβ2-upregulated genes which were mostly associated with biosynthetic pathways including lipid and amino acid metabolism (data not shown).

## DISCUSSION

*S. pseudintermedius* is one of the most common pathogens isolated from the skin of dogs and becoming increasingly prevalent on humans. The incidence of severe and recurrent infections in animals and humans caused by methicillin-resistant *S. pseudintermedius* is increasing and is associated with a predominant clone belonging to the multi-locus sequence type ST71 (Darlow et al., 2017; Perreten et al., 2010; Riegel et al., 2011; Robb et al., 2017; Stegmann et al., 2010; Weese et al., 2009). ST71 isolates exhibit resistance to many classes of antibiotics and represent a considerable therapeutic challenge. Our group has made advancements in the discovery and characterization of skin antimicrobial isolates that are effective against drug-resistant pathogens and can be applied as a novel therapeutic (Cogen et al., 2010; Nakatsuji et al., 2018, 2017; O’Neill et al., 2020). Here we followed a similar discovery pathway, screening for antimicrobial activity in isolates from domestic cats and dogs that could inhibit the growth of *S. pseudintermedius*. We selected an isolate from feline skin that secreted small peptides which were easily extractable by established biochemical methods for an antimicrobial extract preparation. The application of either treatment modality onto infected mouse skin greatly reduced MRSP infection and colonization. These results highlight the clinical potential for transplantation of antimicrobial commensals to treat skin infections that are recalcitrant to classic antibiotic treatment.

There are several reports of the discovery and characterisation of distinct antimicrobial-producing strains within common human bacterial skin species, including *S. epidermidis* (Cogen et al., 2010) *S. hominis* (Nakatsuji et al., 2017), *S. lugdunensis* (Zipperer et al., 2016)*, S. capitis* (O’Neill et al., 2020) and recently *Cutibacterium acnes* (Claesen et al., 2020). Unfortunately, the frequency and abundance of such antimicrobial isolates are found to be significantly reduced during human AD and *S. aureus* colonization (Nakatsuji et al., 2017). Indeed, this phenomenon could also influence canine AD, which shares many of the clinical features of human AD and a corresponding predisposition to *S. pseudintermedius* colonization. We hypothesized that the screening of poorly characterized bacterial species from diverse body sites of healthy animals could be a promising strategy to discover new antimicrobial strains that are effective against zoonotic pathogens. Following the high-throughput antimicrobial screening of a collection of diverse animal-derived staphylococci, we discovered *S. felis* C4, which secretes antimicrobials targeting MRSP and other clinically relevant, drug-resistant gram-positive pathogens. *S. felis* remains poorly characterized in the literature, but is the most frequent species of staphylococci isolated from cats and is susceptible to most antimicrobials (K. Worthing et al., 2018). In this study, twenty-three *S. felis* isolates were screened but only three showed reproducible antimicrobial activity in liquid and agar co-culture, suggesting an uncommon intraspecies trait. Importantly, we demonstrated that topical application of the live *S. felis* C4 organism outcompeted MRSP colonization *in vivo*, likely by the active secretion of its antimicrobials on the skin surface. Indeed, topical application of the sterile extract was similarly effective in reducing CFU counts on mouse skin. Injection of the extract during *S. pseudintermedius* skin infection was also effective in significantly reducing the size of necrotic lesions. These positive *in vivo* findings build upon other reports of the utilization of commensal bacteria as biotherapeutic products to treat skin diseases.

In the treatment of AD, it has been suggested that the topical application of a 5% lysate from the gram-negative bacterium *Vitreoscilla filiformis* can be effective in human and mouse AD models (Gueniche et al., 2008). Another group reported improved outcomes in mice and on humans when another commensal gram-negative bacterium called *Roseomonas mucosa* was applied to their skin. Although gram-negative bacteria do not typically survive on the skin surface, this strain was isolated from the skin of a healthy human volunteer, suggesting transient colonization from environmental exposure (Myles et al., 2019). Our group has sought to isolate potentially protective bacteria from taxa that have evolved the capacity to survive on the skin, thus maximizing the capacity to attack pathogen targets. We have recently shown that *S. aureus* colonization is reduced after transplantation of a human commensal *S. hominis* A9 strain onto AD patient skin in a randomized, double-blind, placebo-controlled trial (Nakatsuji et al., 2017; Nakatsuji et al. in press). This anti-*S. aureus* activity was mediated by several secreted lantibiotics unique to *S. hominis* A9. Typically, lantibiotics and bacteriocins are produced by multimodular enzymes of the nonribosomal peptide synthetases and polyketide synthases (Walsh, 2008). Unfortunately, their complex structure and chemistry are major hurdles to their adoption in the clinic. Furthermore, they are difficult to purify and synthesize, and are unstable and sensitive to proteases, pH, and oxidation (Ross and Vederas, 2011). Alternatively, small α-helical AMPs, which include mammalian cathelicidin (active LL-37) and defensins (Nakatsuji and Gallo, 2012) as well as bacterial PSMs (Cogen et al., 2010; Zeng et al., 2019), could be better candidates.

The family of phenol-soluble modulins which include PSMα, PSMβ, PSMγ, and PSMε, are a class of small, immunomodulatory AMPs found in many staphylococci species. PSMα and PSMγ are classic cytolytic toxins but PSMβ does not exhibit cytotoxic activity against eukaryotic membranes despite a strong affinity to bind and lyse POPC vesicles, which mimic biological membranes (Duong et al., 2012). As a result, the biological function of PSMβ has remained unclear (Cheung et al., 2014). However, several recent studies have reported the antimicrobial activity of PSMβ (Kumar et al., 2017; O’Neill et al., 2020). Here, we detected several PSMβ peptides by MS, that were highly enriched in an antimicrobial HPLC fraction from *S. felis* C4 supernatant. Their activity against the growth of MRSP was validated using synthetic versions of the peptides, but they exhibited less potency than the extract. As such, due to the partially purified nature of the extract, the contribution of other antimicrobials products cannot be ruled out. This could include the AMP PSMα which is known to co-elute with PSMβ by HPLC, but is difficult to detect and not well annotated in bacterial genomes due to its small size (Cheung et al., 2014; Joo et al., 2011). Nevertheless, both fluorescence and electron microscopy of bacteria exposed to the extract showed drastic perturbations of the bacterial cell membrane and cell wall thickening, which is consistent with the membrane-targeting actions of amphipathic AMPs. The concomitant accumulation of bacterial ROS and decrease in ATP production are also consistent with bacterial membrane disruption and increased permeability (Song et al., 2020). Future research efforts will attempt to develop and characterize truncated and mutated versions of *S. felis* PSMs with the goal of enhancing antimicrobial activity for a more simplistic but potentially more powerful therapeutic (Zeng et al., 2019).

In addition to their protection against pathogen colonization, skin commensals play important roles in promoting skin health and immune homeostasis. Although PSMs are common amongst staphylococci, pathogenic *S. aureus* exhibits a preference for PSMα production over PSMβ. In contrast, commensal staphylococci production of PSMβ is prioritized over the more toxic PSMα and PSMγ versions, a feature suspected to be an evolutionary adaptation to stably colonize skin (Otto, 2009; Wang et al., 2007). Naturally, *S. felis* C4 PSMβ and extract treatment of NHEKs yielded minimal evidence of cytotoxicity, whereas PSMα and PSMγ induced extensive cytotoxicity. These smaller PSMs are well characterised toxins that trigger pro-inflammatory responses (Nakamura et al., 2013; Williams et al., 2019). Whereas the larger PSMβ reportedly does not elicit pro-inflammatory activity *in vitro* - a finding that was supported by our data, little is known regarding other potential host responses to PSMβ exposure (Cheung et al., 2014). We speculated whether PSMβ might exhibit anti-inflammatory activities, and in the present context that activity would be therapeutically beneficial. Indeed, when both NHEKs and THP-1 macrophages were treated with *S. felis* PSMs or extract in the presence of TLR agonists, cytokine induction was reduced but most evidently by PSMβ2. RNA-Seq analysis of NHEKs revealed global suppression of inflammatory pathways typically activated by TLR3, in the presence of PSMβ2. Treatment of NHEKs with PSMβ2 alone showed the downregulation of genes associated with IL-1 and bacterial infection, suggesting that PSMβ promotes tolerance to commensal microbes. Here we report that *S. felis* C4 bacterium is an attractive biotherapeutic candidate for skin disease, that could benefit patients due to established low cytotoxicity and its broad-spectrum antimicrobial activity and anti-inflammatory activity.

## MATERIALS AND METHODS

### Bacterial strains and growth conditions

The bacterial strains used in this study were all grown overnight, with the exception of *E. faecium* which was grown for 48 h, in Tryptic Soy Broth (TSB) (Oxoid) with shaking or on agar at 37 °C under static conditions.

### Sample collection

Animal-derived staphylococci samples came from two previously described collections: the first collection consisted of clinical isolates of skin and soft tissue infection from Australian dogs and cats (K. A. Worthing et al., 2018), and the second collection was comprised of staphylococci isolated from the nose, mouth and perineum of healthy dogs and cats in Australia (Ma et al., 2020). All samples had previously been identified by matrix assisted laser desorption ionization-time of flight mass spectrometry (MALDI-TOF), as previously described (K. Worthing et al., 2018), and the ST71 MRSP isolate had been characterized by whole genome sequencing (K. A. Worthing et al., 2018). Two human derived antimicrobial skin commensal isolates were used as positive controls: *S. hominis* A9 (Nakatsuji et al., 2017) and *S. capitis* E12 (O’Neill et al., 2020) and a non-antimicrobial *S. aureus* 113 isolate served as a negative control.

### *In vitro* antimicrobial screen

For the initial staphylococci screen, single clone-derived cultures of animal-derived staphylococci were used as competitor isolates against the growth of methicillin-resistant *S. pseudintermedius* ST71. Each pure culture, including positive and negative control strains, were first streaked onto 3% TSB agar plates and a single colony was transferred to 1 ml of TSB in a deep 96 well plate (Thermo). The CoNS plate was sealed with sterile Aeraseal film (Sigma, St. Louis, MO) and cultured at 37°C overnight with shaking at 250 rpm. Bacterial growth was evaluated by measuring OD600 with only bacteria that grew to a density (OD600 > 6.0) used for subsequent analysis. To measure antimicrobial activity in the secreted supernatant, the animal-derived staphylococci supernatant from overnight cultures were harvested and centrifuged through several 96-well 0.2 μm sterile filter plates (Corning). Next, 1×10^5^ CFU of *S. pseudintermedius* ST71 was inoculated into 150 μl of 100%, 50% or 25% sterile supernatant supplemented with fresh 3% TSB and grown on a plate shaker for 18 h at 30°C. To measure antimicrobial activity from the live agar co-culture assays, 20 μl of overnight *S. pseudintermedius* ST71 culture was first inoculated into 45°C molten TSB and immobilized after pouring and cooling into square petri dishes with grids. Overnight cultures of animal-derived staphylococci were centrifuged to pellet the bacteria, washed 2X with PBS and resuspended in fresh TSB. The culture (10 μl) was inoculated onto a 13 mm grid of the *S. pseudintermedius* agar plates and cultured overnight at 30°C. The resulting zones of inhibition from antimicrobial isolates were imaged using the camera feature on an iPhone 12.

### Extraction and purification of antimicrobials from bacterial supernatant

Supernatant from overnight cultures of selected human and animal-derived staphylococci were first sterilized by filtration through a 0.22-μm Millipore filter. Activity was precipitated by ammonium sulfate (75% saturation) for 1 h, under constant rotation followed by centrifugation at 4,000 x g for 45 min and re-suspension of the pellet in dH_2_O. To test stability, the precipitate was boiled at 95°C for 30 minutes or stored in a sterile eppendorf tube at room temperate for 1 week. Antimicrobial activity was measured by radial diffusion assay. Sterile supernatant of S. *felis* strains were subject to n-Butanol extraction and purification as previously reported (Joo and Otto, 2014). Briefly, in each tube 10 ml of butanol was added to 30 ml of supernatant and incubated at 37°C for 2 h under constant rotation. The tubes were then set aside for several mins until the butanol phase settled. After centrifugation at 2000 x g for 5 min, the upper butanol phase was collected and lyophilized in a SpeedVac vacuum concentrator. The lyophilized extract was resuspended and concentrated to 10 mg/ml in DMSO.

### Determination of minimum inhibitory concentration (MIC)

MIC values were determined using a broth micro dilution method. Bacterial cells were grown to mid-late log phase, to an OD600 nm value of roughly 1.0 for each bacterial strain and then normalized to 1×10^7^ CFU/mL. The PSM peptides or butanol extracts were dissolved in DMSO to a stock concentration of 10 mg/ml. The stock concentrations of antibiotics that were water-soluble were prepared with H_2_O or 100% ethanol if water-insoluble. The 1×10^7^ CFU/ml bacterial cultures (10 μl) were aliquoted into 96-well microtiter plates and mixed with 95 μL of media with or without 2-fold dilutions of the conditioned supernatant, PSM peptides, butanol extracts or antibiotics and incubated for 16–18 h at 30°C with shaking at 250 rpm. Growth inhibition was determined by measuring the OD600 nm readings of each well using a microplate reader (SpectraMax iD3, Molecular Devices). The MIC of each bacterial strain was determined by the lowest peptide concentration that inhibited more than 80% bacterial growth.

### Crystal Violet staining for biofilm disruption

Overnight culture of *S. pseudintermedius* ST71 was diluted in fresh TSB to 1×10^7^ CFU/ml by OD600 reading. A total of 100 μl of bacteria was transferred to a flat-bottom 96 well plate and incubated at 37°C without shaking for 4 h to initiate biofilm formation. Next, the supernatants were removed by washing the plates three times with 200 μl of dH_2_O. Subsequently, 150 μl of *S. felis* C4 supernatant, extract, or PSMβ2 peptide at various concentrations, or TSB negative control, was added to the biofilm for periods between 2 – 24 h. After incubation, the supernatant was gently removed, and the biofilm was washed three times with dH_2_O followed by air drying. Next, 150 μl of 0.1% crystal violet (CV) solution was added to all wells containing biofilm. After 20 mins of incubation with CV dye, the excess CV was removed and each well was washed twice with dH_2_O. Fixed CV dye was released from the biofilm by 70% ethanol, and absorbance was measured at 595 nm.

### HPLC purification and peptide synthesis

First step HPLC purification was carried out with 1 mg of *S. felis* C4 supernatant loaded onto a Capcell Pak® C8 column (5 mm, 300 A°, 4.6 mm 250 mm) (Shiseido, Tokyo, Japan) using a linear acetonitrile gradient from 10% to 60% in 0.1% (v/v) trifluoroacetic acid at a flow rate of 1.0 ml/min. The resulting fractions were lyophilized, then resuspended in water, and antimicrobial activity assessed by liquid culture assay. Up to five sequential purifications were carried out with each antimicrobial fraction pooled together for the second HPLC purification. A linear gradient of acetonitrile from 25% to 50% was used for the second purification. PSM peptides were synthesized with N-terminal formulation to at least 95% purity by a commercial vendor (LifeTein LLC, Somerset, NJ).

### Silver Stain and acetone precipitation of antimicrobial fractions

Twenty μg of protein from sources including the antimicrobial HPLC fractions, butanol extracts and crude supernatant were loaded onto a Novex 16% Tricine gel and subjected to SDS-PAGE. Silver staining and de-staining of the protein gels were performed according to the manufacturer’s instructions (Thermo Pierce Silver Stain Kit). A previously published protocol for acetone extraction of AMPs from SDS gels was used (Burgess, 2009). Briefly, a sterile razor blade was used to excise gel slices according to protein size. The gel slices were cut into small pieces and immersed in dH_2_0 for 4 h, with regular vortexing to elute proteins. The eluted protein was mixed with four volumes of ice-cold acetone for 1 h at −20°C. The samples were centrifuged at 16,000 x g for 15 mins at 4°C. The supernatant was removed and lyophilized (acetone-soluble fraction) and the resulting pellet air dried briefly and resuspended in dH_2_0 (acetone-insoluble fraction). The antimicrobial activity of both fractions was tested by radial diffusion agar assay against *S. pseudintermedius* ST71.

### Immunoblot

NHEK cells were treated with Poly I:C (0.4 μg/ml), PSMβ2 (10 μg/ml) or Poly I:C and PSMβ2 combined for 15 min, 30 min or 60 min and cells lysed in complete RIPA buffer supplemented with 1× protease and phosphatase inhibitor cocktail (Life Technologies, USA). The lysate was centrifuged at 4°C, at 13,000 rpm for 20 min and the total cytoplasmic supernatant fraction was kept at −80 °C, until future use. The total protein amount was quantified for each treatment using the Pierce™ BCA Protein Assay Kit according to manufacturer’s instructions. Fifteen μg of total protein was loaded onto a 4–20% Mini-PROTEAN TGX gel (Bio-Rad), then transferred to a polyvinylidene difluoride (PVDF) membrane and probed with the following primary antibodies: P-TBK1/NAK (D52C2), TBK1/NAK (D1B4), P-IRF-3 (S396), IRF-3 (D83B9), COX IV (3E11). IRDye conjugated anti-rabbit and anti-mouse secondary antibodies (IRDye800CW; Licor, USA) were used. The images were acquired on an Odyssey CLx Imaging System (Licor, USA).

### Mass spectrometry

Four fractions of interest (ranging from 10-20 μg/mL) were dried under vacuum and resuspended in 15 μL of 5% acetonitrile with 5% formic acid. Next, individual LC-MS experiments were conducted on 6 μL of each sample through 85 minutes of data acquisition on an Orbitrap Fusion (Thermo Fisher Scientific) mass spectrometer with an in-line Easy-nLC 1000 (Thermo Fisher Scientific). A home-pulled and packed 30 cm column was triple-packed with 0.5 cm, 0.5 cm and 30 cm of 5 μm C4, 3 μm C18, and 1.8 μm C18 respectively and heated to 60 °C for use as the analytical column. Peptides were first loaded at 500 bar which was followed by a chromatography gradient ranging from 6 to 25% acetonitrile over 70 minutes followed by a 5-minute gradient to 100% acetonitrile, which was held for 10 minutes. Electrospray ionization was performed by applying 2000V through a stainless-steel T-junction connecting the analytical column and Easy-nLC system. Each sample was followed by four washes starting with a gradient from 3 to 100% acetonitrile over 15 minutes with an additional 10 minutes at 100% acetonitrile. An m/z range of 375-1500 was scanned for peptides with charge states between 2-6. Centroided data was used for quantitation of peaks. Acquisition was run in a data-dependent positive ion mode. Raw spectra was searched in Proteome Discoverer Version 2.1 against 6-frame translated databases based of a uniprot reference database for *Staphylococcus felis* ATCC 49168 (Uniprot proteome UP000243559, accessed 06/26/2019) as well as in-house sequencing of *S. felis* C4. Data were searched using the Sequest algorithm (Eng et al., 1994) using a reverse database approach to control peptide and protein false discoveries to 1% (Elias and Gygi, 2007). No enzyme was specified in the search and a minimum peptide length was set to 6 amino acids. Search parameters included a precursor mass tolerance of 50 ppm and fragment mass tolerance of 0.6 Da and variable oxidation for modifications.

### Whole genome sequencing of *S. felis* C4

DNA was extracted from *S. felis* C4 using the UltraClean™ Microbial DNA Isolation Kit (MoBio) according to the manufacturer’s instructions. The library was prepared using Nextera DNA Flex library preparation kit according to the manufacturer’s instructions (Illumina, San Diego, CA). The library was diluted to 1.0nM, then sequenced for 300 cycles using the Illumina NovaSeq system to generate 150bp paired-end reads with 794x coverage that was reduced to 100x coverage for read mapping. Fastq files from *S. felis* C4 were trimmed using Trimmomatic (Bolger et al., 2014), then assembled using SPAdes Genome Assembler v.3.14.1. The *S. felis* C4 genome was annotated using the RAST tool kit via the Pathosystems Resource Integration Center (PATRIC) database (Wattam et al., 2014) and genes encoding PSMβ were identified by BLASTn.

### ATP determination

The intracellular ATP levels of *S. pseudintermedius* ST71 treated with *S. felis* C4 butanol extract were measured following the manufacturer’s instructions (ReadiUse™ Rapid Luminometric ATP Assay Kit). Briefly, an overnight *S. pseudintermedius* ST71 culture was sub-cultured to an OD of 0.5 at 37°C. The bacteria were pelleted, washed with fresh TSB and incubated with various concentrations (0-320 μg/ml) of *S. felis* C4 butanol extract for 1 h. The bacterial cultures were centrifuged at 10,000 x g for 5 min at 4°C. The bacterial pellet was lysed by lysozyme and centrifuged. The bacterial supernatant was mixed with an equal volume of detecting solution in a 96 well plate and incubated at room temperature for 20 min. ATP luminescence was read using a SpectraMax iD3 (Molecular Devices).

### Reactive Oxygen Species (ROS) measurement

The levels of reactive oxygen species (ROS) in *S. pseudintermedius* ST71 that was treated with *S. felis* C4 butanol extract were measured with 2’,7’-dichlorofluorescein diacetate (DCFDA) following the manufacturer’s instructions (DCFDA/H2DCFDA Abcam cellular ROS assay kit). Briefly, an overnight culture of *S. pseudintermedius* ST71 was sub-cultured to an OD of 0.5 at 37°C. The bacteria were pelleted and re-suspended in fresh TSB. DCFDA was added to a final concentration of 20 μM to the bacterial culture incubated with various concentrations of extract (0-320 μg/ml) at 37°C for 1 hour. Fluorescence intensity was immediately measured at an excitation wavelength of 488 nm and an emission wavelength of 525 nm using a SpectraMax iD3 (Molecular Devices).

### Bacterial viability assay

Dead or damaged bacteria induced by *S. felis* C4 extract were evaluated using the LIVE/DEAD® *Bac*Light™ Bacterial Viability Kit (Invitrogen, catalogue no. L7007). An overnight *S. pseudintermedius* ST71 culture was washed with fresh TSB and OD adjusted to 0.5 under treatment with different concentrations of extract (0, 0.2, 8, and 32 μg/ml). After incubation for 1 h, the bacteria were harvested, washed and resuspended in 1X PBS. SYTO9 (1.67mmol1–1, 1.5 μl) and PI (10mmol1–1, 1.5 μl) were added to each sample to a final volume of 1 ml and incubated at room temperature in the dark for 15 min. Fluorescent images of stained bacteria were obtained using a EVOS M5000 fluorescent microscope.

### Transmission electron microscopy

*S. pseudintermedius* ST71 cell pellets were immersed in modified Karnovsky’s fixative (2% glutaraldehyde and 2% paraformaldehyde in 0.10 M sodium cacodylate buffer, pH 7.4) for at least 4 h and further postfixed in 1% osmium tetroxide in 0.1 M cacodylate buffer for 1 hr on ice. The cells were stained all at once with 2% uranyl acetate for 1 hr on ice, then dehydrated in a graded series of ethanol (50-100%) while remaining on ice. The cells were washed with 100% ethanol and washed twice with acetone (10 min each) and embedded with Durcupan. Sections were cut at 60 nm on a Leica UCT ultramicrotome, and picked up on 300 mesh copper grids. Sections were post-stained with 2% uranyl acetate for 5 minutes and Sato’s lead stain for 1 minute. Grids were viewed using a JEOL JEM-1400Plus (JEOL, Peabody, MA) transmission electron microscope and photographed using a Gatan OneView 4K digital camera (Gatan, Pleasanton, CA).

### Mouse skin colonization and infection with *S. pseudintermedius*

#### Mouse skin colonization

All experiments involving live animal work were performed in accordance with the approval of the University of California, San Diego Institutional Animal Care and Use Guidelines (protocol no. S09074). For mouse skin challenge experiments involving *S. felis* strains and *S. pseudintermedius* ST71, the dorsal skin of hairless age-matched 8-10 week-old SKH1 mice (n=2, per treatment) were scrubbed with alcohol wipes and 5×10^6^/cm^2^ or 5×10^7^/cm^2^ CFU of overnight cultured *S. felis* C4, *S. felis* ATCC 49168 or *S. pseudintermedius* ST71 was inoculated onto 1 x 1 cm sterile gauze pads, which were placed onto the dorsal skin and secured with wound dressing film (Tegaderm [3M]) film for 72 h. For experiments involving live *S. felis* C4 bacteria or extract topical treatment, the dorsal skin of age-matched 8-10 week-old C57BL/6 mice (n=4, per treatment) was shaved and depilated by using Nair cream followed by removal with alcohol wipes. The skin was allowed to recover from hair removal for at least 24 h before the application of bacteria. Prior to bacterial challenge, the dorsal skin was tape-stripped and *S pseudintermedius* ST71 agar disks (3% tryptic soy broth [TSB], 2% agar; diameter 8 mm) containing 5 x 10^7^ CFU was applied to the skin for 48 h, as previously described (Nakatsuji et al., 2016). The dorsal skin was covered with Tegaderm and a bandage was applied to hold the agar disk or gauze in place for the duration of the treatment. The bandage, Tegaderm and agar disk were removed and 5×10^7^/cm^2^ CFU of overnight cultured *S. felis* C4, or 150 μl of extract (10 mg/ml) or 3% TSB control, was inoculated onto 2 x 2 cm sterile gauze pads and applied to the infected site every 24 h for 72 h. After the treatment of mouse skin with live bacteria or extract, the dressing film and gauze pad were removed, and surface bacteria were collected using a swab soaked in TSB-glycerol solution. The swab head was then placed in 1 mL of TSB-glycerol solution, vortexed (30 seconds), serial-diluted, and plated onto Baird Parker agar plates supplemented with egg yolk tellurite for enumeration of coagulase-positive staphylococci (Carter, 1960) or mannitol salt agar plates for enumeration of all surface staphylococci (Parisi and Hamory, 1986).

#### Mouse infection and dermonecrosis model

The day prior to bacterial infection, the dorsal skin of age-matched 8-10 week old C57BL/6 mice (n=5, per treatment) was shaved and depilated by using Nair cream followed by removal with alcohol wipes. A 50 μl inoculum suspension containing 1 x 10^7^ CFU of late log phase *S. pseudintermedius* ST71 in PBS was intradermally injected into the dorsal skin using 0.3 mL/31-gauge insulin syringe (BD, Franklin Lakes, NJ). At 1 h post-infection, a 50 μl suspension of the S. *felis* C4 extract (250 μg, at a concentration of 5 mg/ml in 25% DMSO) or a control suspension of 1 X PBS in 25% DMSO was injected twice in two separate skin sites directly adjacent to the bacterial injection site. Body weights of the mice were measured before and after infection every day for 14 days. To determine lesion size, a ruler was positioned adjacent to the mouse skin lesions and digital photos were taken daily with a Kodak PIXPRO Astro Zoom AZ421 and analyzed via ImageJ software (National Institutes of Health Research Services Branch, Bethesda, MD, USA). Lesion size in mm^2^ was measured by calculating the length x width.

### Quantitative real-time PCR

mRNA transcript abundance was measured by qPCR in NHEKs stimulated with or without TLR2/6 agonist MALP-2 (200 ng/ml) or TLR3 agonist Poly I:C (0.4 μg/ml) in the presence or absence of *S. felis* C4 extract, PSMβ2 or PSMβ3 (10 μg/ml) or DMSO control (0.1%) at 4 h post-treatment. RNA was extracted from NHEK cells using Pure Link RNA isolation kit (Life Technologies, USA) according to manufacturer’s instructions. RNA was quantified on a Nanodrop 2000/200c spectrophotometer (Thermo Fisher, USA). Purified RNA (500 μg) was used to synthesize cDNA using the iScript™ cDNA Synthesis Kit (Bio-Rad, USA). Pre Developed Taqman® (Thermo Fisher, USA) and SYBR-Green gene expression assays (Integrated DNA Technologies, USA) were used to evaluate mRNA transcript levels.

### RNA Sequencing

NHEK cells were treated with DMSO (0.1%) control, PSMβ2 (10 μg/ml), Poly I:C (0.4 μg/ml) or PSMβ2 and Poly I:C combined, all in triplicate wells, for 4 h or 24 h in and RNA was extracted using the PureLink RNA mini kit and triplicate samples were pooled together for each treatment. Isolated RNA was submitted to the UCSD IGM Genomics Center for RNA-sequencing performed on a high-output run V4 platform (Illumina, USA) with a single read 100 cycle runs. Data alignment was performed using Partek® Flow® genomic analysis software (Partek, USA) with Tophat2 (version 2.0.8) Gene ontology (GO) enrichment analysis was performed on differentially regulated genes (≥1.5-fold) using DAVID 6.8.

### Statistical analysis

Significant differences between the means of the different treatments were evaluated using GraphPad Prism version 7.03 (GraphPad Software, Inc., La Jolla, CA). Either unpaired, two-tailed Student’s *t* test or one-way analysis of variance (ANOVA) followed by Dunnett’s or Turkey’s multiple comparisons test were used for statistical analysis and indicated in the respective figure legends. Differences were considered statistically significant with a *p* value of <0.05.

## ACKNOWLEDGEMENTS

The authors would like to acknowledge the participants for their assistance in this project. We thank Ying Jones of the UCSD/CMM electron microscopy facility for TEM sample preparation and Timothy Meerloo for imaging assistance. The EM facility is supported by NIH equipment grant 1S10OD023527. We thank Nina J. Gao for providing strains. We thank Jamie Boehmer for her very helpful edits. We also thank Gayathri Kalla, Joe Pogliano, Kit Pogliano and Gemma Ma.

## FUNDING

This publication includes data generated at the UC San Diego IGM Genomics Center utilizing an Illumina NovaSeq 6000 that was purchased with funding from a National Institutes of Health SIG grant (#S10 OD026929). R.H.M. was supported through a UCSD training grant from the NIH/NIDDK Gastroenterology Training Program (T32 DK007202).

## CONFLICT OF INTEREST

R.L.G. is a co-founder, scientific advisor, consultant and has equity in MatriSys Biosciences and is a consultant, receives income and has equity in Sente. K.W. and R.L.G are co-inventors of technology described in this report that has been disclosed to the University of California San Diego.

## SUPPLEMENTARY FIGURES

**Suppl. Figure 1.**
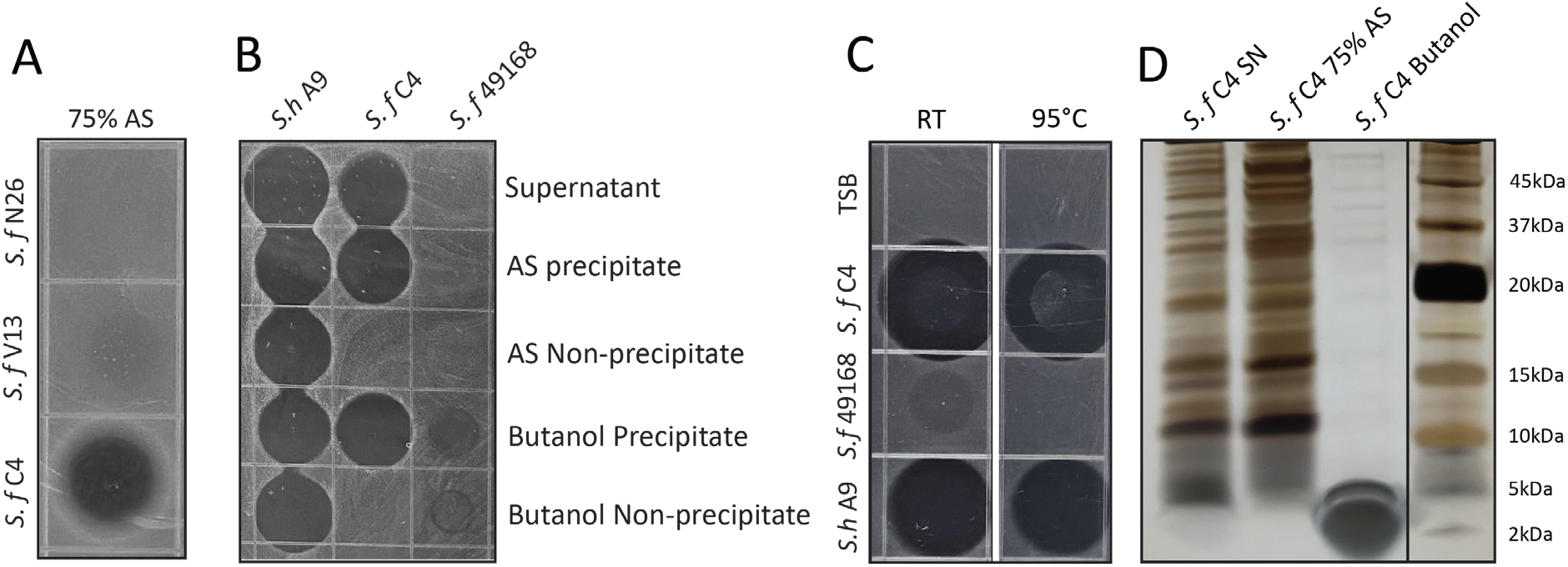
Generation of a partially purified antimicrobial extract from *S. felis* C4. **(A)** Supernatant of indicated antimicrobial *S. felis* strains (*S.f* C4, N26, V13) were incubated with 75% ammonium sulfate (AS) and the resulting precipitate was inoculated onto agar containing *S. pseudintermedius* ST71, to determine antimicrobial activity. **(B)** Supernatant of *S. felis* C4, positive-control *S. hominis* A9, or negative-control *S. felis* ATCC 49168 was extracted in 75% AS or 25% n-butanol and activity of the precipitate and non-precipitate fractions were assayed against *S. pseudintermedius* ST71. **(C)** Supernatant of *S. felis* C4 (*S. f* C4), *S. hominis* A9 (*S. h* A9), *S. capitis* E12 (*S. c* E12) or TSB alone, were maintained at room temperature (RT) for 1 week or boiled at 95°C, for 30 mins and activity determined by inoculation directly onto agar containing *S. pseudintermedius* ST71. **(D)** Total protein silver stain of *S. felis* C4 supernatant before precipitation or after precipitation in 75% AS or 25% butanol.

**Suppl. Figure 2.**
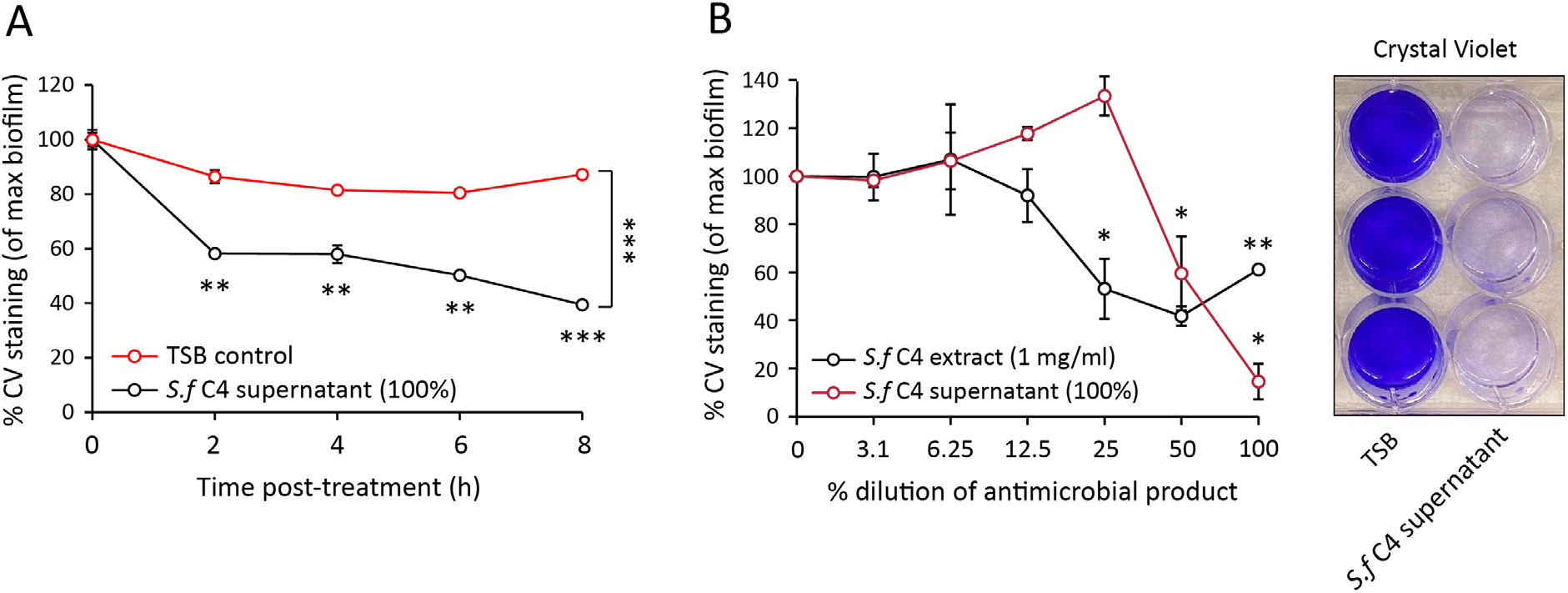
*S. felis* C4 supernatant and extract disrupt *S. pseudintermedius* biofilm. **(A)** Decrease in crystal violet staining of a 4 h preformed *S. pseudintermedius* ST71 biofilm during 8 h incubation with 100% *S. felis* C4 sterile supernatant. **(B)** Decrease in crystal violet staining of a 4 h preformed *S. pseudintermedius* ST71 biofilm after 24 h incubation with serially diluted *S. felis* C4 sterile supernatant (100%) or *S. felis* C4 extract (1 mg/ml). Right inset is a representative image of crystal violet staining after incubation with 100% *S. felis* C4 supernatant. (**A-B**) Error bars indicate SEM. Representative of two separate experiments. A two-tailed, unpaired Student’s t test was performed. *p* values: \**p* < 0.05; \*\**p* < 0.01; \*\*\**p* < 0.001.

**Suppl. Figure 3.**
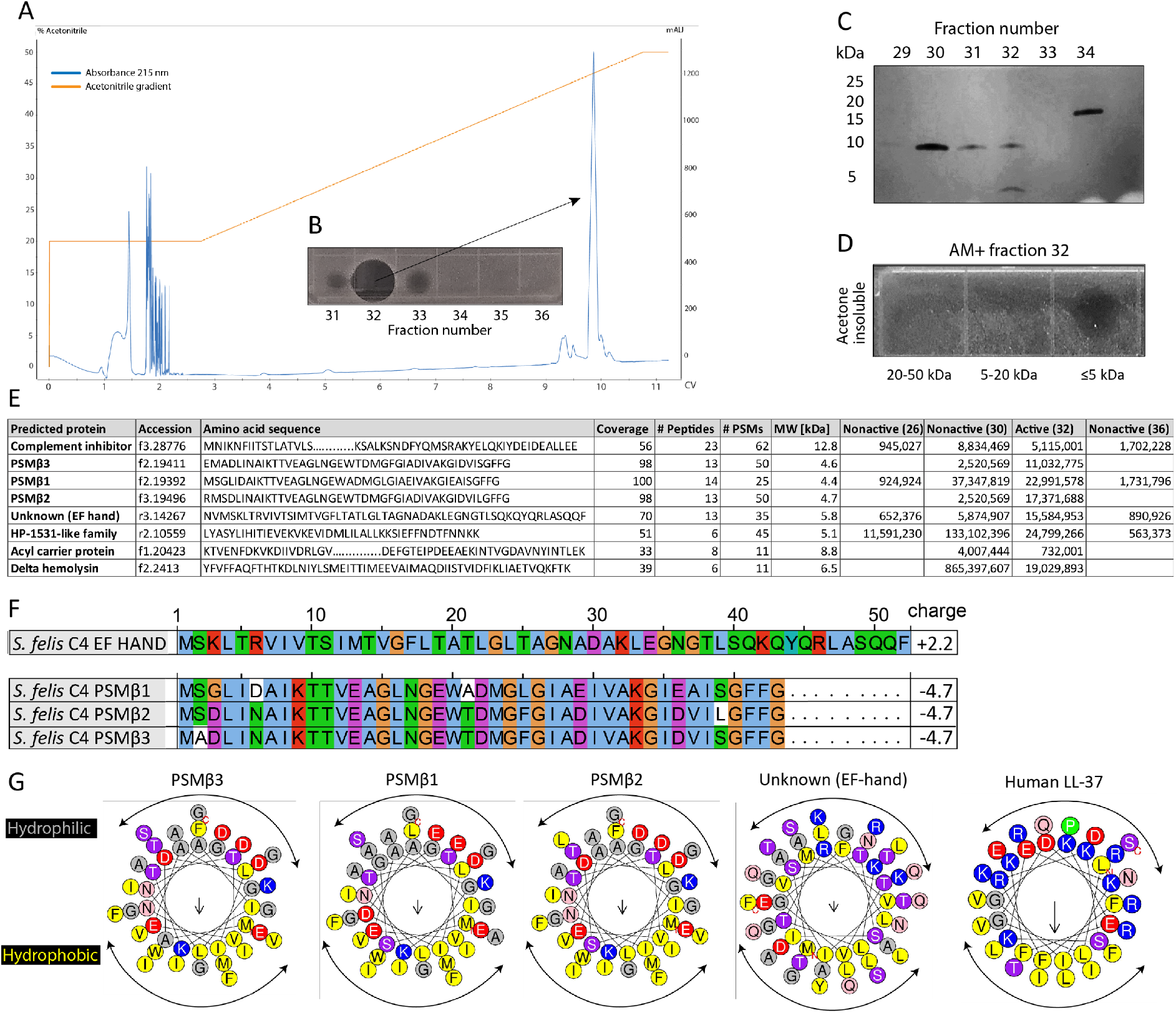
HPLC purification yields antimicrobial fraction from *S. felis* C4 supernatant. **(A)** Reverse-phase high-performance liquid chromatography (HPLC) elution profile from sterile supernatant of *S. felis* C4 strain loaded onto a C8 column. **(B)** Inset image of antimicrobial activity exhibited by fraction 32 against *S. pseudintermedius* ST71 corresponding to the indicated peak. **(C)** Silver stain of total protein content in the different fractions indicated. **(D)** Radial diffusion assay of antimicrobial activity of the AM+ fraction 32 after extraction and acetone precipitation of proteins within different sized silver stain gel fragments. **(E)** Mass spectrometry (MS) table of the top 8 peptide hits obtained from HPLC fractions that were active (fraction 32) or inactive (26, 30, 36) against *S. pseudintermedius* ST71. **(F)** ClustalW multiple amino acid sequence alignment of all three *S. felis* C4 genetically-encoded PSMβ peptides with predicted net charge at pH 7.4 (Prot pi) and amino acid sequence of a EF-hand domain-containing peptide with unknown function. **(G)** Alpha helical wheel plots of *S. felis* C4 PSMβ1-3, EF-hand domain peptide and human LL-37 peptide, indicating conserved α-helical, amphipathic-like structures with indicated hydrophobic yellow residues confined to one side (indicated by arrow) and grey hydrophilic residues on the opposing side.

**Suppl. Figure 4.**
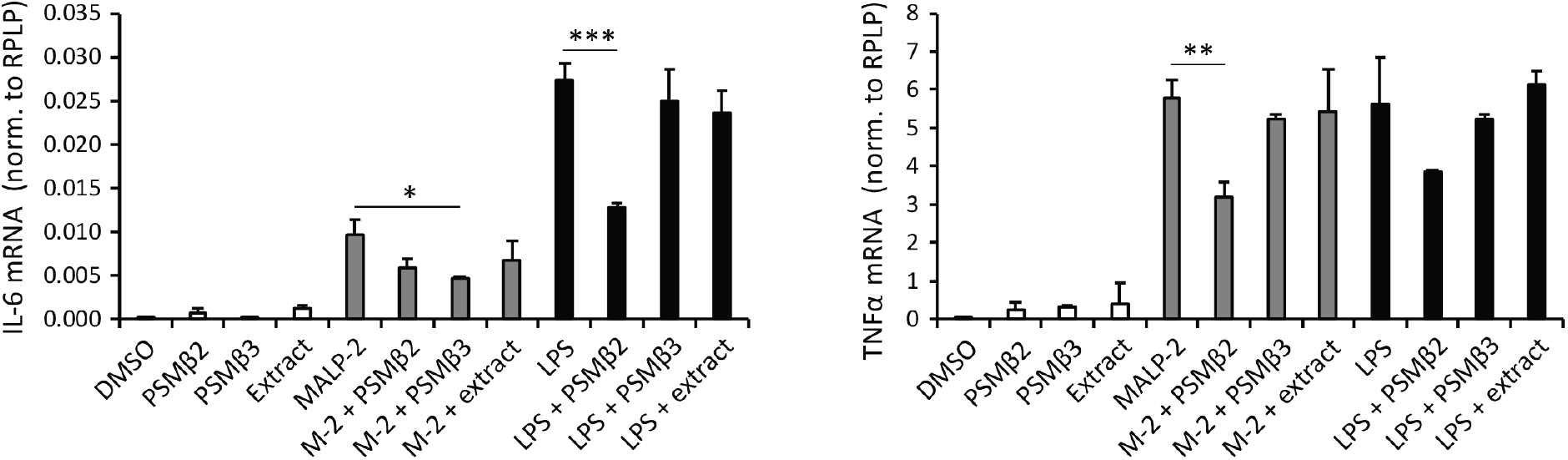
PSMβ2 reduces TLR2- and TLR4-stimulated transcripts in THP-1 macrophages. Transcript abundance of inflammatory cytokines IL-6 and TNFα in THP-1 cells stimulated with or without TLR2/6 agonist MALP-2 (200 ng/ml) or TLR4 agonist LPS (1 μg/ml) in the presence or absence of *S. felis* C4 extract, PSMβ2 or PSMβ3 (10 μg/ml) or DMSO control (0.1%) 4 h post-treatment. Error bars indicate SEM. One-way ANOVA with multiple corrections (Turkey correction) was performed.*p* values: * p< 0.05; **, p<0.01; *** p<0.001.

**Suppl. Figure 5.**
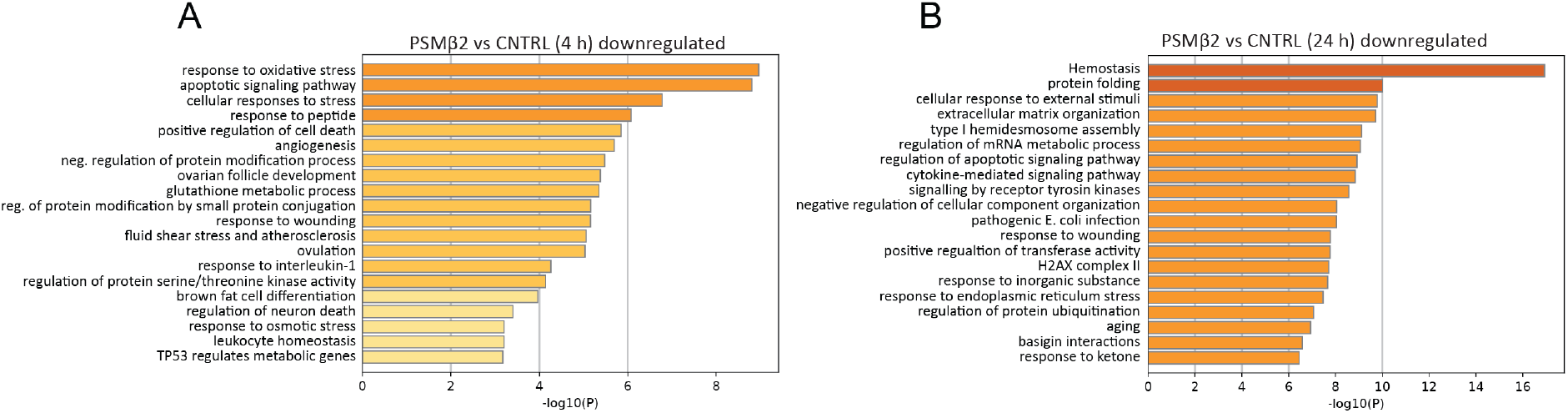
PSMβ2 supresses TLR3-mediated cellular activation and downregulates responses to bacterial pathogens. **(A-B)** GO pathway analysis of genes down-regulated in NHEKs at 4 h **(A)** and 24 h **(B)** after treatment with PSMβ2 versus DMSO control.

## REFERENCES

Banovic F, Olivry T, Bäumer W, Paps J, Stahl J, Rogers A, Jacob M. 2018. Diluted sodium hypochlorite (bleach) in dogs: antiseptic efficacy, local tolerability and in vitro effect on skin barrier function and inflammation. Vet Dermatol 29:5–6. doi:10.1111/vde.12487

Bolger AM, Lohse M, Usadel B. 2014. Trimmomatic: A flexible trimmer for Illumina sequence data. Bioinformatics 30:2114–2120. doi:10.1093/bioinformatics/btu170

Börjesson S, Gómez-Sanz E, Ekström K, Torres C, Grönlund U. 2015. *Staphylococcus pseudintermedius* can be misdiagnosed as *Staphylococcus aureus* in humans with dog bite wounds. Eur J Clin Microbiol Infect Dis 34:839–844. doi:10.1007/s10096-014-2300-y

Botelho D, Wall MJ, Vieira DB, Fitzsimmons S, Liu F, Doucette A. 2010. Top-down and bottom-up proteomics of sds-containing solutions following mass-based separation. J Proteome Res 9:2863–2870. doi:10.1021/pr900949p

Bradley CW, Morris DO, Rankin SC, Cain CL, Misic AM, Houser T, Mauldin EA, Grice EA. 2016. Longitudinal evaluation of the skin microbiome and association with microenvironment and treatment in canine atopic dermatitis. J Invest Dermatol 136:1182–1190. doi:10.1016/j.jid.2016.01.023

Burgess RR. 2009. Chapter 32 Elution of proteins from gelsmethods in enzymology. Academic Press Inc. pp. 565–572. doi:10.1016/S0076-6879(09)63032-9

Carter CH. 1960. Egg yolk agar for isolation of coagulase-positive staphylococci. J Bacteriol 79.

Chazin WJ. 2011. Relating form and function of EF-hand calcium binding proteins. Acc Chem Res 44:171–179. doi:10.1021/ar100110d

Cheung GYC, Joo HS, Chatterjee SS, Otto M. 2014. Phenol-soluble modulins - critical determinants of staphylococcal virulence. FEMS Microbiol Rev. doi:10.1111/1574-6976.12057

Chopra R, Vakharia PP, Sacotte R, Silverberg JI. 2017. Efficacy of bleach baths in reducing severity of atopic dermatitis: A systematic review and meta-analysis. Ann Allergy, Asthma Immunol 119:435–440. doi:10.1016/j.anai.2017.08.289

Claesen J, Spagnolo JB, Ramos SF, Kurita KL, Byrd AL, Aksenov AA, Melnik A V., Wong WR, Wang S, Hernandez RD, Donia MS, Dorrestein PC, Kong HH, Segre JA, Linington RG, Fischbach MA, Lemon KP. 2020. A *Cutibacterium acnes* antibiotic modulates human skin microbiota composition in hair follicles. Sci Transl Med 12. doi:10.1126/scitranslmed.aay5445

Cogen AL, Yamasaki K, Muto J, Sanchez KM, Alexander LC, Tanios J, Lai Y, Kim JE, Nizet V, Gallo RL. 2010. *Staphylococcus epidermidis* antimicrobial δ-toxin (phenol-soluble modulin-γ) cooperates with host antimicrobial peptides to kill group A Streptococcus. PLoS One 5. doi:10.1371/journal.pone.0008557

Da F, Joo H-S, Cheung GYC, Villaruz AE, Rohde H, Luo X, Otto M. 2017. Phenol-soluble modulin toxins of *Staphylococcus haemolyticus*. Front Cell Infect Microbiol 7:206. doi:10.3389/fcimb.2017.00206

Darlow CA, Paidakakos N, Sikander M, Atkins B. 2017. A spinal infection with *Staphylococcus pseudintermedius*. BMJ Case Rep 2017. doi:10.1136/bcr-2017-221260

Di Domenico EG, Cavallo I, Bordignon V, Prignano G, Sperduti I, Gurtner A, Trento E, Toma L, Pimpinelli F, Capitanio B, Ensoli F. 2018. Inflammatory cytokines and biofilm production sustain *Staphylococcus aureus* outgrowth and persistence: A pivotal interplay in the pathogenesis of Atopic Dermatitis. Sci Rep 8. doi:10.1038/s41598-018-27421-1

Duong AC, Cheung GYC, Otto M. 2012. Interaction of phenol-soluble modulins with phosphatidylcholine vesicles. Pathogens 1:3–11. doi:10.3390/pathogens1010003

Elias JE, Gygi SP. 2007. Target-decoy search strategy for increased confidence in large-scale protein identifications by mass spectrometry. Nat Methods 4:207–214. doi:10.1038/nmeth1019

Eng JK, McCormack AL, Yates JR. 1994. An approach to correlate tandem mass spectral data of peptides with amino acid sequences in a protein database. J Am Soc Mass Spectrom 5:976–989. doi:10.1016/1044-0305(94)80016-2

Fazakerley J, Nuttall T, Sales D, Schmidt V, Carter SD, Hart CA, McEwan NA. 2009. Staphylococcal colonization of mucosal and lesional skin sites in atopic and healthy dogs. Vet Dermatol 20:179–184. doi:10.1111/j.1365-3164.2009.00745.x

Ference EH, Danielian A, Kim HW, Yoo F, Kuan EC, Suh JD. 2019. Zoonotic *Staphylococcus pseudintermedius* sinonasal infections: risk factors and resistance patterns. Int Forum Allergy Rhinol 9:724–729. doi:10.1002/alr.22329

Frana TS, Beahm AR, Hanson BM, Kinyon JM, Layman LL, Karriker LA, Ramirez A, Smith TC. 2013. Isolation and characterization of methicillin-resistant *Staphylococcus aureus* from pork farms and visiting veterinary students. PLoS One 8:e53738. doi:10.1371/journal.pone.0053738

Garbacz K, Zarnowska S, Piechowicz L, Haras K. 2013. Pathogenicity potential of *Staphylococcus pseudintermedius* strains isolated from canine carriers and from dogs with infection signs. Virulence. doi:10.4161/viru.23526

Grice EA, Segre JA. 2011. The skin microbiome. Nat Rev Microbiol 9:244. doi:10.1038/nrmicro2537

Gueniche A, Knaudt B, Schuck E, Volz T, Bastien P, Martin R, Röcken M, Breton L, Biedermann T. 2008. Effects of nonpathogenic gram-negative bacterium *Vitreoscilla filiformis* lysate on atopic dermatitis: A prospective, randomized, double-blind, placebo-controlled clinical study. Br J Dermatol 159:1357–1363. doi:10.1111/j.1365-2133.2008.08836.x

Joo HS, Cheung GYC, Otto M. 2011. Antimicrobial activity of community-associated methicillin-resistant *Staphylococcus aureus* is caused by phenol-soluble modulin derivatives. J Biol Chem 286:8933–8940. doi:10.1074/jbc.M111.221382

Joo HS, Otto M. 2014. The isolation and analysis of phenol-soluble modulins of *Staphylococcus epidermidis*. Methods Mol Biol 1106:93–100. doi:10.1007/978-1-62703-736-5_7

Kong HH, Oh J, Deming C, Conlan S, Grice EA, Beatson MA, Nomicos E, Polley EC, Komarow HD, Mullikin J, Thomas J, Blakesley R, Young A, Chu G, Ramsahoye C, Lovett S, Han J, Legaspi R, Sison C, Montemayor C, Gregory M, Hargrove A, Johnson T, Riebow N, Schmidt B, Novotny B, Gupta J, Benjamin B, Brooks S, Coleman H, Ho SL, Schandler K, Stantripop M, Maduro Q, Bouffard G, Dekhtyar M, Guan X, Masiello C, Maskeri B, McDowell J, Park M, Vemulapalli M, Murray PR, Turner ML, Segre JA. 2012. Temporal shifts in the skin microbiome associated with disease flares and treatment in children with atopic dermatitis. Genome Res 22:850–859. doi:10.1101/gr.131029.111

Kumar R, Jangir PK, Das J, Taneja B, Sharma R. 2017. Genome analysis of *Staphylococcus capitis* TE8 reveals repertoire of antimicrobial peptides and adaptation strategies for growth on human skin. Sci Rep 7. doi:10.1038/s41598-017-11020-7

Kwiecinski J, Kahlmeter G, Jin T. 2015. Biofilm formation by *staphylococcus aureus* isolates from skin and soft tissue infections. Curr Microbiol 70:698–703. doi:10.1007/s00284-014-0770-x

Lai PS, Allen JG, Hutchinson DS, Ajami NJ, Petrosino JF, Winters T, Hug C, Wartenberg GR, Vallarino J, Christiani DC. 2017. Impact of environmental microbiota on human microbiota of workers in academic mouse research facilities: An observational study. PLoS One 12:e0180969. doi:10.1371/journal.pone.0180969

Ma GC, Worthing KA, Ward MP, Norris JM. 2020. Commensal Staphylococci Including Methicillin-resistant *Staphylococcus aureus* from dogs and cats in remote New South Wales, Australia. Microb Ecol 79:164–174. doi:10.1007/s00248-019-01382-y

Mandhane PJ, Sears MR, Poulton R, Greene JM, Lou WYW, Taylor DR, Hancox RJ. 2009. Cats and dogs and the risk of atopy in childhood and adulthood. J Allergy Clin Immunol 124:745–750.e4. doi:10.1016/j.jaci.2009.06.038

Marsella R, Girolomoni G. 2009. Canine models of atopic dermatitis: A useful tool with untapped potential. J Invest Dermatol. doi:10.1038/jid.2009.98

Myles IA, Williams KW, Reckhow JD, Jammeh ML, Pincus NB, Sastalla I, Saleem D, Stone KD, Datta SK. 2019. Transplantation of human skin microbiota in models of atopic dermatitis. JCI Insight 1. doi:10.1172/jci.insight.86955

Nakamura Y, Oscherwitz J, Cease KB, Chan SM, Muñoz-Planillo R, Hasegawa M, Villaruz AE, Cheung GYC, McGavin MJ, Travers JB, Otto M, Inohara N, Núñez G. 2013. Staphylococcus δ-toxin induces allergic skin disease by activating mast cells. Nature 503:397–401. doi:10.1038/nature12655

Nakatsuji T, Chen TH, Butcher AM, Trzoss LL, Nam SJ, Shirakawa KT, Zhou W, Oh J, Otto M, Fenical W, Gallo RL. 2018. A commensal strain of *Staphylococcus epidermidis* protects against skin neoplasia. Sci Adv. doi:10.1126/sciadv.aao4502

Nakatsuji T, Chen TH, Narala S, Chun KA, Two AM, Yun T, Shafiq F, Kotol PF, Bouslimani A, Melnik A V., Latif H, Kim JN, Lockhart A, Artis K, David G, Taylor P, Streib J, Dorrestein PC, Grier A, Gill SR, Zengler K, Hata TR, Leung DYM, Gallo RL. 2017. Antimicrobials from human skin commensal bacteria protect against *Staphylococcus aureus* and are deficient in atopic dermatitis. Sci Transl Med. doi:10.1126/scitranslmed.aah4680

Nakatsuji T, Chen TH, Two AM, Chun KA, Narala S, Geha RS, Hata TR, Gallo RL. 2016. *Staphylococcus aureus* exploits epidermal barrier defects in atopic dermatitis to trigger cytokine expression. J Invest Dermatol. doi:10.1016/j.jid.2016.05.127

Nakatsuji T, Gallo RL. 2019. The role of the skin microbiome in atopic dermatitis. Ann Allergy, Asthma Immunol. doi:10.1016/j.anai.2018.12.003

Nakatsuji T, Gallo RL. 2012. Antimicrobial peptides: Old molecules with new ideas. J Invest Dermatol. doi:10.1038/jid.2011.387

O’Neill AMAM, Nakatsuji T, Hayachi A, Williams MRMR, Mills RHRH, Gonzalez DJDJ, Gallo RLRL. 2020. Identification of a human skin commensal bacterium that selectively kills *Cutibacterium acnes*. J Invest Dermatol 140:1619–1628.e2. doi:10.1016/j.jid.2019.12.026

Older CE, Hoffmann AR, Hoover K, Banovic F. 2020. Characterization of cutaneous bacterial microbiota from superficial pyoderma forms in atopic dogs. Pathogens 9:1–12. doi:10.3390/pathogens9080638

Otto M. 2009. Staphylococcus epidermidis - The “accidental” pathogen. Nat Rev Microbiol. doi:10.1038/nrmicro2182

Parisi JT, Hamory BH. 1986. Simplified method for the isolation, identification, and characterization of Staphylococcus epidermidis in epidemiologic studies. Diagn Microbiol Infect Dis 4:29–35. doi:10.1016/0732-8893(86)90053-2

Perreten V, Kadlec K, Schwarz S, Andersson UG, Finn M, Greko C, Moodley A, Kania SA, Frank LA, Bemis DA, Franco A, Iurescia M, Battisti A, Duim B, Wagenaar JA, van Duijkeren E, Weese JS, Fitzgerald JR, Rossano A, Guardabassi L. 2010. Clonal spread of methicillin-resistant *Staphylococcus pseudintermedius* in Europe and North America: An international multicentre study. J Antimicrob Chemother 65:1145–1154. doi:10.1093/jac/dkq078

Riegel P, Jesel-Morel L, Laventie B, Boisset S, Vandenesch F, Prévost G. 2011. Coagulase-positive *Staphylococcus pseudintermedius* from animals causing human endocarditis. Int J Med Microbiol 301:237–239. doi:10.1016/j.ijmm.2010.09.001

Robb AR, Wright ED, Foster AME, Walker R, Malone C. 2017. Skin infection caused by a novel strain of *Staphylococcus pseudintermedius* in a Siberian husky dog owner. JMM Case Reports 4. doi:10.1099/jmmcr.0.005087

Ross AA, Müller KM, Scott Weese J, Neufeld JD. 2018. Comprehensive skin microbiome analysis reveals the uniqueness of human skin and evidence for phylosymbiosis within the class Mammalia. Proc Natl Acad Sci U S A 115:E5786–E5795. doi:10.1073/pnas.1801302115

Ross AC, Vederas JC. 2011. Fundamental functionality: Recent developments in understanding the structure-activity relationships of lantibiotic peptides. J Antibiot (Tokyo), doi:10.1038/ja.2010.136

Sanford JA, Gallo RL. 2013. Functions of the skin microbiota in health and disease. Semin Immunol. doi:10.1016/j.smim.2013.09.005

Sawada Y, Tong Y, Barangi M, Hata T, Williams MR, Nakatsuji T, Gallo RL. 2019. Dilute bleach baths used for treatment of atopic dermatitis are not antimicrobial in vitro. J Allergy Clin Immunol. doi:10.1016/j.jaci.2019.01.009

Silverberg JI. 2019. Comorbidities and the impact of atopic dermatitis. Ann Allergy, Asthma Immunol 123:144–151. doi:10.1016/j.anai.2019.04.020

Singh A, Walker M, Rousseau J, Weese JS. 2013. Characterization of the biofilm forming ability of *Staphylococcus pseudintermedius* from dogs. BMC Vet Res 9. doi:10.1186/1746-6148-9-93

Somayaji R, Priyantha MAR, Rubin JE, Church D. 2016. Human infections due to *Staphylococcus pseudintermedius*, an emerging zoonosis of canine origin: report of 24 cases. Diagn Microbiol Infect Dis 85:471–476. doi:10.1016/j.diagmicrobio.2016.05.008

Song J, Lauber C, Costello EK, Lozupone CA, Humphrey G, Berg-Lyons D, Caporaso JG, Knights D, Clemente JC, Nakielny S, Gordon JI, Fierer N, Knight R. 2013. Cohabiting family members share microbiota with one another and with their dogs. elife.elifesciences.org Song al eLife 2:458. doi:10.7554/eLife.00458

Song M, Liu Y, Huang X, Ding S, Wang Y, Shen J, Zhu K. 2020. A broad-spectrum antibiotic adjuvant reverses multidrug-resistant Gram-negative pathogens. Nat Microbiol 5:1040–1050. doi:10.1038/s41564-020-0723-z

Stegmann R, Burnens A, Maranta CA, Perreten V. 2010. Human infection associated with methicillin-resistant *Staphylococcus pseudintermedius* ST71. J Antimicrob Chemother 65:2047–2048. doi:10.1093/jac/dkq241

Walsh CT. 2008. The chemical versatility of natural-product assembly lines. Acc Chem Res 41:4–10. doi:10.1021/ar7000414

Wang R, Braughton KR, Kretschmer D, Bach THL, Queck SY, Li M, Kennedy AD, Dorward DW, Klebanoff SJ, Peschel A, DeLeo FR, Otto M. 2007. Identification of novel cytolytic peptides as key virulence determinants for community-associated MRSA. Nat Med 13:1510–1514. doi:10.1038/nm1656

Wattam AR, Abraham D, Dalay O, Disz TL, Driscoll T, Gabbard JL, Gillespie JJ, Gough R, Hix D, Kenyon R, MacHi D, Mao C, Nordberg EK, Olson R, Overbeek R, Pusch GD, Shukla M, Schulman J, Stevens RL, Sullivan DE, Vonstein V, Warren A, Will R, Wilson MJC, Yoo HS, Zhang C, Zhang Y, Sobral BW. 2014. PATRIC, the bacterial bioinformatics database and analysis resource. Nucleic Acids Res 42:D581. doi:10.1093/nar/gkt1099

Weese JS, Poma R, James F, Buenviaje G, Foster R, Slavic D. 2009. Case report. Can Vet J 50:655–656.

Williams MR, Costa SK, Zaramela LS, Khalil S, Todd DA, Winter HL, Sanford JA, O’Neill AM, Liggins MC, Nakatsuji T, Cech NB, Cheung AL, Zengler K, Horswill AR, Gallo RL. 2019. Quorum sensing between bacterial species on the skin protects against epidermal injury in atopic dermatitis. Sci Transl Med. doi:10.1126/scitranslmed.aat8329

Worthing K, Pang S, Trott DJ, Abraham S, Coombs GW, Jordan D, McIntyre L, Davies MR, Norris J. 2018. Characterisation of *Staphylococcus felis* isolated from cats using whole genome sequencing. Vet Microbiol 222:98–104. doi:10.1016/j.vetmic.2018.07.002

Worthing KA, Abraham S, Coombs GW, Pang S, Saputra S, Jordan D, Trott DJ, Norris JM. 2018. Clonal diversity and geographic distribution of methicillin-resistant *Staphylococcus pseudintermedius* from Australian animals: Discovery of novel sequence types. Vet Microbiol 213:58–65. doi:10.1016/j.vetmic.2017.11.018

Zeng P, Xu C, Cheng Q, Liu J, Gao W, Yang X, Wong KY, Chen S, Chan KF. 2019. Phenol-soluble-modulin-inspired amphipathic peptides have bactericidal activity against multidrug-resistant bacteria. ChemMedChem 14:1547–1559. doi:10.1002/cmdc.201900364

Zhang Y, Bottinelli D, Lisacek F, Luban J, Strambio-De-Castillia C, Varesio E, Hopfgartner G. 2015. Optimization of human dendritic cell sample preparation for mass spectrometry-based proteomic studies. Anal Biochem 484: 40–50. doi:10.1016/j.ab.2015.05.007

Zipperer A, Konnerth MC, Laux C, Berscheid A, Janek D, Weidenmaier C, Burian M, Schilling NA, Slavetinsky C, Marschal M, Willmann M, Kalbacher H, Schittek B, Brötz-Oesterhelt H, Grond S, Peschel A, Krismer B. 2016. Human commensals producing a novel antibiotic impair pathogen colonization. Nature 535:511–516. doi:10.1038/nature18634

